# Altered Extracellular Matrix Structure and Elevated Stiffness in a Brain Organoid Model for Disease

**DOI:** 10.1101/2024.01.09.574777

**Authors:** Maayan Karlinski Zur, Bidisha Bhattacharya, Sivan Ben Dror, Inna Solomonov, Alon Savidor, Tamar Sapir, Talia Harris, Tsviya Olender, Irit Sagi, Rita Schmidt, J. M. Schwarz, Amnon Buxboim, Orly Reiner

## Abstract

The viscoelasticity of tissues impacts their shape, as well as the growth and differentiation of their cells. Nevertheless, little is known about changes in viscoelastic properties during brain malformations. Lissencephaly is a severe malformation of cortical development caused by LIS1 mutations, which results in a lack of cortical convolutions. Here, we show that human-derived brain organoids with *LIS1* mutation are stiffer than control ones at multiple developmental times. This stiffening is accompanied by abnormal ECM expression and organization, as well as elevated water content, as measured by diffusion-weighted MRI. Proteolytic cleavage of ECM components by short-term treatment with the catalytic subunit of MMP9 reduced the stiffening and water diffusion levels of mutated organoids to control levels. Finally, based on the molecular and rheological properties obtained, we generated a computational microstructure mechanical model that can successfully predict mechanical changes that follow differential ECM localization and integrity in the developing brain. Overall, our study reveals that LIS1 is essential for the expression and organization of ECM proteins during brain development, and its mutation leads to a substantial viscoelastic change. To our knowledge, this is the first study to elucidate how tissue mechanics change in disease states using human brain organoids.

## Introduction

Tissue mechanics is crucial in shaping tissue growth, function, and disease progression. However, this field is understudied in human brain developmental diseases due to limited access to human brain tissues and suboptimal animal models ^1–6^. It has been suggested that the tissue composition, the dynamic cellular processes that occur in the developing brain, and tissue mechanics play a role in shaping brain structure ^7–19^. The organization, shape, and amount of structural extracellular matrix (ECM) in the tissue provide tissues with their mechanical forces, thereby affecting cellular behavior during and after development ^20, 21^. ECM has been proposed to be a crucial element in the structural organization of the developing brain and the formation of human brain folds ^22, 23^. Lissencephaly, characterized by the absence of cortical convolutions, provides insights into human brain fold formation. LIS1 mutations, prevalent in lissencephaly patients, impact the scaffold protein LIS1, affecting cytoplasmic dynein, RNA interactions, splicing, and gene transcription ^24–26^. Studying this disease in mouse models has been useful but limited since their cortex naturally lacks convolutions. The mutant mice exhibited deficits in neuronal migration and hippocampal pathology ^27–35^. Here, we used human pluripotent stem cell-derived organoids to study how biomechanical changes are involved in cortical malformation development. We found that brain organoids mutated for *LIS1* are stiffer than control organoids and unraveled a substantial ECM disorganization phenotype in the disease. Using an interdisciplinary approach, we applied data obtained from rheological tests, MRI, and ECM composition and structure characterization to develop a computational model. This model successfully predicts mechanical changes associated with differential ECM localization and integrity in the developing brain.

## Results

### Mutant *LIS1* brain organoids are stiffer

To investigate whether the cortical structural abnormalities observed in cases of lissencephalic pathologies are linked to mechanical abnormalities, we performed rheological tests on cortical organoids (CorticOs). These organoids were generated from two types of cell lines: control human embryonic stem cells (hESCs) and isogenic lines with a *LIS1* heterozygous mutation introduced using CRISPR/Cas9 genome editing techniques ^16^.

CorticOs were generated using a protocol for generating self-organizing cortical tissue (Supplementary **Fig. S1a**). A series of characterizations on different days indicated that the early ectoderm-like organoids were expressing different neural progenitors on days 9 and 18. By day 60 corticOs contained post-mitotic neurons and astrocytes (Supplementary **Fig. S1b-e**), and to enrich for basal radial glial progenitors, thought to play an important role in the developing human cortex^36^, we added hLIF from day 35. Basal radial glial progenitors were detected in 96 days old corticOs (supplementary **Fig. S1f-g**).

Changes in mechanics can alter the structure, development, and function of cells that make up a tissue, such as the brain ^45^. To determine the mechanical effects of *LIS1* mutations, we employed micropipette aspiration (MPA) rheology and performed creep test measurements of brain organoids from *LIS1^+/−^* and control organoids at multiple developmental ages (days 9, 18, 35, and 70). The aspiration dynamics of the organoids into the pipette under a constant negative pressure were recorded and analyzed **(Fig. 1a)**. All organoid measurements independent of developmental age shared a characteristic response to the applied load: organoids stretched elastically the moment suction was applied, followed by a gradual aspiration into the pipette over five to ten seconds, and approached a steady-state finite deformation **(Supplementary Fig. S2a)**.

**Fig. 1:**
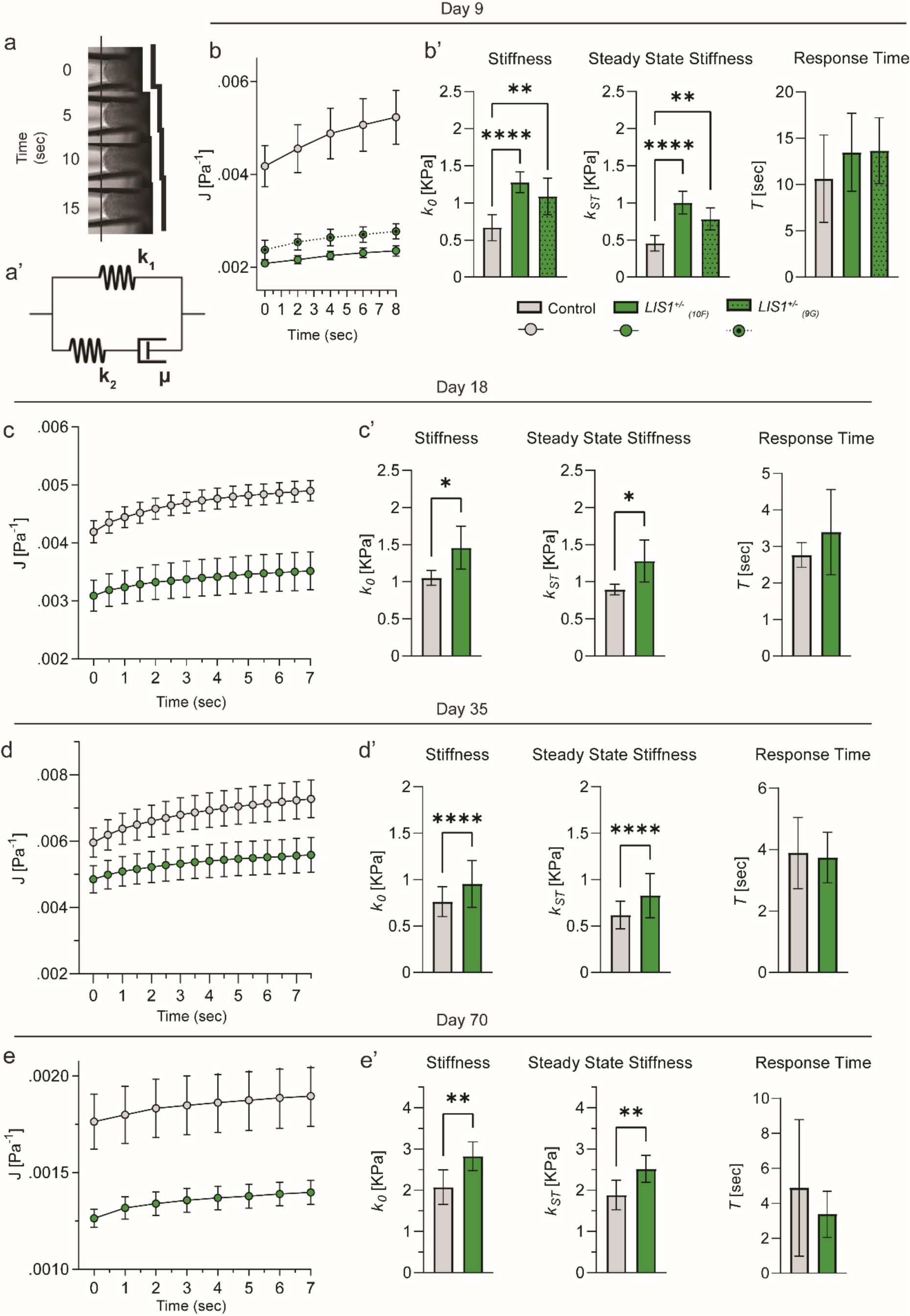
Organoids are solid-like viscoelastic that stiffen by ECM deposition downstream of *LIS1* mutations. **a**, Under constant suction pressure, the organoids are continuously aspirated into the pipette and gradually approach a steady-state deformation (**a**). The organoids’ creep compliance is well-fitted by the standard linear solid (SLS) viscoelastic model (**a’**). **b-e,** Averaged creep compliance measurements (symbols) are fitted by the SLS model (curves) at the specified conditions. Symbols and error bars correspond to the mean and standard error of the mean. (**b’-e’**) SLS fits the instantaneous (k_0_) and steady-state (k_st_) stiffness and response time (τ) are plotted. The analysis included the following number of organoids, each measured separately and fitted independently: (**b)** Day-9, N_WT_ = 6, N_10F_ = 8, N_9G_ = 7. (**c)** Day-18, N_WT_ = 4, N_10F_ = 5. **d)** Day-35, N_WT_ = 8, N_10F_ = 8. (**e)** Day-70, N_WT_ = 7, N_10F_ = 9. Statistical significance is evaluated via a two-way ANOVA test.

The mechanical behaviour of organoids was analysed using the standard linear solid (SLS) model. In its Maxwell representation, it consists of an elastic element (spring *k_1_*) that is connected in parallel to a second elastic element (spring *k_2_*) positioned in series with a viscous element (dashpot *µ*) **(Fig. 1a’)**. We calculated the creep compliance function, J(t), to measure time-dependent deformability^49^, using parameters like the aspirated fraction length, pipette radius, applied pressure, and a geometrical factor^50^. The organoid mechanics were quantitatively characterized by fitting the SLS creep compliance function, considering factors like instantaneous stiffness, steady-state stiffness, and the transition time scale. This model showed high accuracy in representing organoid behaviour, as evidenced by the high R-square values in our fits (See **Methods** for a full description of the model and its calculation).

CorticOs stiffness, as estimated by *k_0_* and *k_st_*, ranged over hundreds of pascals, indicating that it is as soft as cream cheese **(Fig. 1b-e)** ^51^. Notably, this range aligns with the lower spectrum of brain tissue stiffness, which typically spans from 0.1 to 2 kilopascals ^52^. With time, the CorticOs stiffen, likely due to continuous ECM deposition and/or fibrillation. We found that *LIS1^+/−^* mutations increased the stiffness of corticOs as early as 9 days after their aggregation, and that they remain stiffer than controls up to the latest tested time point, day 70. However, no significant difference in the response time is observed **(Fig. 1b’-e’)**. These findings indicate that, when subjected to a physiologically relevant load across multicellular length scales, the CorticOs exhibit characteristics of a material with strain-stiffening behaviour. This stiffening tendency becomes more pronounced with progressing developmental stages. Overall, our data suggests that changes to the biomechanics of *LIS1*^+/−^ organoids appear early in development and continues over a considerable period.

### Cortical *LIS1^+/−^* organoids express abnormal ECM and more Lamin A

To delineate the molecular changes that are associated with the stiffening of the *LIS1^+/−^* cortices, we analyzed the proteomic signature of both control and *LIS1^+/−^* corticOs **(Supplementary Table S1a)**. Previous studies indicated that basal radial glial cells may be involved in cortical gyrification (review ^37^). Therefore, we have chosen to conduct analysis in 105-day-old corticOs, following the appearance and expansion of that progenitor population **(Supplementary Fig. S1f-g)**. On day 105, there were 253 DE proteins between the *LIS1^+/−^* and control corticOs **(Fig. 2a)**. The top affected pathway identified by www.geneanalytics.com ^38^ was the super-path “chromatin regulation/acetylation”, followed by the “Super path: ECM organization” **(Supplementary Table S1b)**.

**Fig. 2:**
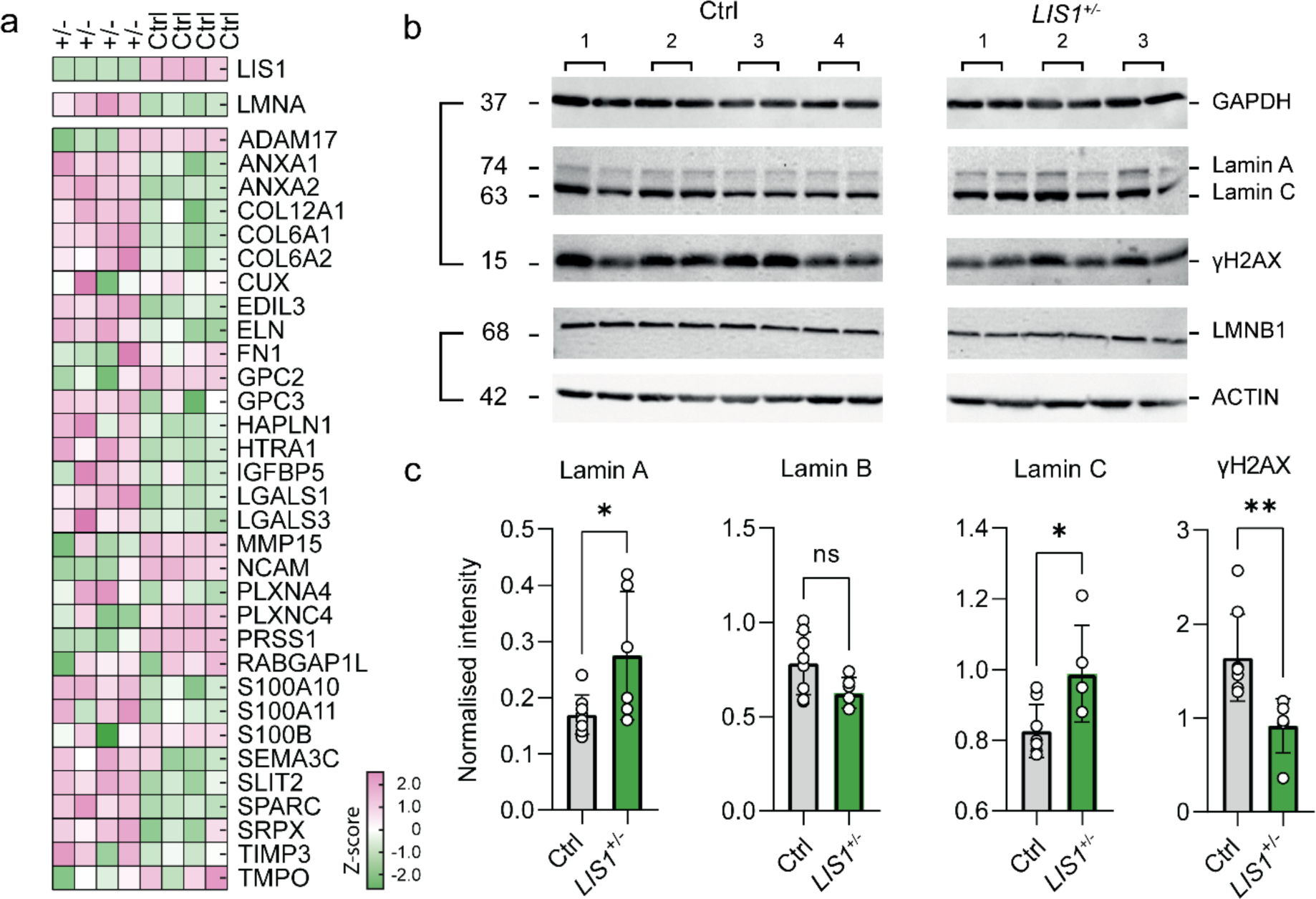
Matrisome composition of CorticO. **a,** Heatmap of top DE matrisomal proteins in *LIS1^+/−^* and control 105 days old cortical organoids. Cell color express normalized reads following logarithmic transformation and Z-score normalization **b,** Western blot analysis indicated the elevation of Lamin A/C and the reduction in gH2AX in 105 days old *LIS1^+/−^* CorticOs. **c**, Quantification of E (Two-tailed unpaired t-test, α = 0.05, * means p-value < 0.1 and ** means p-value < 0.01).

In line with the observed stiffening of the corticOs in the MPA procedure, we also observed increased expression of the LMNA protein in the *LIS1^+/−^* corticOs **(Fig. 2a)**. Lamins are intermediate filament proteins of the nucleus that provide structural stability. Lamin A expression is strongly correlated with tissue stiffness, whereas other members of the nuclear lamina family, Lamin B, and Lamin C, are not affected by the physical properties of the cells ^39^. The roles of LMNA are not strictly structural as it also affects chromatin organization, gene regulation, cell differentiation, and signaling pathways including the Wnt/β-catenin pathway, TGFβ, and Notch ^40^.

The 30% increase in LMNA protein levels indicated by the proteomics was recapitulated by western blot analysis, whereas the levels of Lamin B were unchanged **(Fig. 2b-c)**. It was further hypothesized that the rise in Lamin A would reduce the double-strand breaks in the tissue due to an increased nuclear protective shield. This was supported by the Western blot quantification, which showed a significant reduction in the double strand breaks marker γ-H2AX in the *LIS1*^+/−^ 105 days old corticOs. Overall, the increased levels of Lamin A, together with the changes observed in ECM-related proteins, suggest that mutation in LIS1 results in adverse biomechanical abnormalities to the brain organoids.

In addition, RNA-seq analysis of the same 105 days old corticOs cohort revealed a total of 2,061 DE genes between the *LIS1^+/−^* and control samples **(Supplementary Fig. S3a, Supplementary Table S2)**. Pathway analysis of DE genes indicated that here, too, the collagen-associated pathway was the most affected GO-term in the mutation **(Supplementary Fig. S3b)**. Accordingly, genes involved in generating, processing, or regulating collagens were affected in *LIS1^+/−^*CorticOs, such as several non-fibrillar collagens, including *COL12A1*, *COL14A1*, *COL16A1*, and *COL24A1*. Other collagen coding genes were downregulated in the mutation, including different α-chains of collagen type IV and *COL5A3*. ^41^ ^38, 42, 43^

### Hippocampal *LIS1* organoids exhibit increased ECM

Mutations in *LIS1* have a substantial impact on the organization and structure of the cerebral cortex and some abnormalities have been noted in the hippocampus (review ^44^). The *Lis1* mouse models display pronounced hippocampal abnormalities, but the human hippocampal pathophysiology has not been extensively characterized ^27–35^. We generated control and *LIS1^+/−^* mutated hippocampal organoids (hippOs). The hippOs were generated by exposing the 18-day-old aggregates to a temporal BMP4 and WNT activation (**Supplementary Fig. S4a**) ^45^. After 70 days, the tissue contained hippocampal-like ZBTB20^+^ and LEF1^+^ cells, SOX2^+^, PAX6^+^ and HOPX^+^ progenitors, NeuN^+^ and MAP2^+^ neurons, and GFAP^+^ astrocytes (**Supplementary Fig. S4b-h**).

Using Mass spectrometry, we explored differences in the protein content of control and mutant hippOs ^16^ **(Supplementary Fig. S4i)**. A total of 5,068 proteins were identified and quantified **(Supplementary Table S3)**, of which 1,178 were DE in the mutation, based on the threshold criteria of *p* < 0.05, Log2Fold change ≥|0.5|, and >1 peptide per protein. The proteome confirmed a considerable reduction of the mutant LIS1 protein and revealed that *LIS1^+/−^* organoids were highly enriched with ECM-related proteins **(Supplementary Fig. S4i)**. Differentially expressed (DE) ECM proteins included an increased level of structural proteins in the mutant, including several collagen types. These excessive collagens undergo efficient hydroxylation in the mutant organoids, as assessed by the post-translational modifications (PTM) analysis extracted from the mass spectrometry data (**Supplementary Fig. S4j, Supplementary Table S4**). Increased levels of the enzymes involved in these modifications (P3H1-3 and PLOD-3) in the mutant samples further support this. Co- and post-translational hydroxylation of proline residues are essential for maintaining the stability of the triple helical collagen structure, which plays a crucial role in brain development by influencing the properties of the ECM, as well as interactions between cells and their nearby collagens, thus impacting cell behaviour ^46–48^.

In addition to the enrichment in structural proteins, we also observed increased levels of ECM remodelling proteins, such as MMP-14 and LOX, as well as serine protease inhibitors such as SERPINA3, SERPINE2, SERPINB6, and others. These observations were also supported by Gene Set Enrichment Analysis (GSEA), which highlighted the enrichment of genes associated with ECM, collagen fibril organization, and more (**Supplementary Fig. S4k)**. Overall, these findings suggest that brain organoid models representing the human hippocampus also demonstrated pathology linked to the mutation. This aligns with the limited human data available and is in line with observations from previous mouse models.

### Rescue treatment with MMP9 softened the *LIS1* mutated organoids

The stiffening effect of *LIS1^+/−^* mutations can be explained by the altered regulation of ECM secretion and remodeling, as demonstrated by our mass-spec data. To examine whether the stiffness derived from excessive structural ECM proteins, we treated day18 CorticOs with the catalytic subunit of MMP9. This zinc-dependent endopeptidase can cleave multiple ECM and non-ECM fibers broadly expressed in the brain ^49, 50^. The activity of the MMP9 catalytic domain was tested first at different concentrations by an ELISA activity assay **(Supplementary Fig. S5)**. CorticOs were then immersed with 500 µM MMP9 catalytic domain for 10 min and submitted for MPA rheology **(Fig. 3a)**. Despite the proteolytic treatment, the organoids maintained an SLS-like aspiration dynamics and the creep compliance functions indicated a significant increase in deformability **(Fig. 3b)**. Indeed, MMP proteolytic treatment decreased both *k*_0_and *k*_st_ by ∼15% in non-mutated organoids and ∼50% in *LIS1^+/−^* mutated organoids relative to nontreated organoids. Notably, the response time *τ* remained invariant to ECM digestion **(Fig. 3b’)**.

**Fig. 3:**
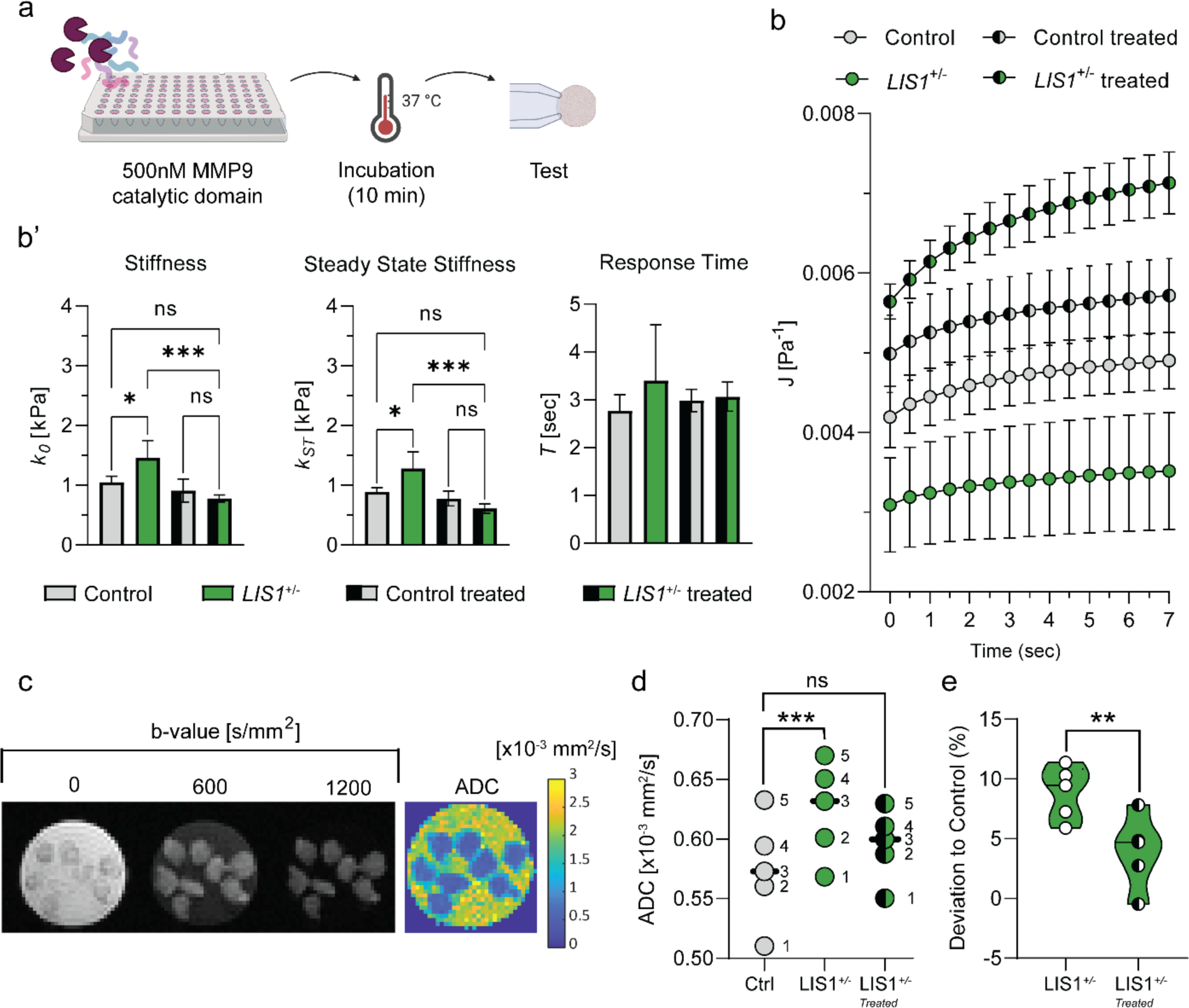
The effects of *LIS1^+/−^*mutation and ECM proteolysis on CorticOs mechanics and structural organization. **a,** Day-18 control and *LIS1^+/−^* mutated organoids were treated by 500 nM catalytic domain for 10 min at 37^0^C before being submitted for MPA creep test measurement and MRI imaging. **b,** Similar to the nontreated organoids, the creep compliance of MMP9-treated organoids of both genotypes were fitted to the linear viscoelastic SLS model with a high goodness of fit (R-square (R-square > 0.99). Symbols and error bars correspond to mean and SEM across N_ctrl-MMP9_ = 5, N_mutant-MMP9_ = 4 organoids. **b’**, Averages ± SEM of the fitted SLS viscoelastic elements k_0_, k_st_, and τ show greater softening of the mutated CorticOs by ECM proteolysis. **c,** A representative DW-MRI of eight *LIS1*^+/−^ corticOs (left) and the calculated ADC map. One central slice is shown. Three out of six b-values (the degree of diffusion weighting) are shown for a central slice. **d,** Estimated maximal likelihood position of the ADC (Apparent Diffusion Coefficient) distributions of control, *LIS1*^+/−^ and MMP9-treated *LIS1^+/−^* corticOs show the effect of altered genotype rescued by ECM proteolysis. Error bars correspond to STD across N = 5 consecutively repeated scans. **e,** the relative deviations of the ADC values of *LIS1*^+/−^ and MMP9-treated *LIS1*^+/−^ corticOs from control organoids are plotted. The potential impact of longitudinal drift was eliminated by calculating each scan’s relative deviation, which was then averaged.

The effects of *LIS1^+/−^* mutations and ECM proteolysis on organoid mechanics are likely associated with structural remodelling. To test this directly, we employed Diffusion-Weighted Magnetic Resonance Imaging (DW-MRI) to compare differences in the structural organization of day18 control, *LIS1*^+/−^, and MMP-treated *LIS1*^+/−^ CorticOs. Notably, DW-MRI was previously shown to provide noninvasive means for identifying changes in ECM organization ^51^, extracellular free water component ^52^, stiffness, and other structural parameters ^53^. To examine the sensitivity of DW-MRI, groups of control, *LIS1*^+/−^, and MMP-treated *LIS1*^+/−^ containing multiple CorticOs were placed in separate wells and scanned **(Fig. 3c-left)**. Taking advantage of the ultra-high magnetic field (15 T MRI scanner), we reached a high imaging resolution of 100×100 μm^2^ in-plane and 200 μm slice thickness voxels. Apparent diffusion coefficient (ADC) maps were calculated **(Fig. 3c-right)**, and the maximal likelihood position was estimated for each group **(Fig. 3d)**. To improve statistical significance, five consecutive scans were performed and analyzed by a matched *t-test;* the matching was effective (P<0.0001). The ADC of *LIS1^+/−^* organoids exhibited a statistically significant 8.9 ± 2.2 % increase in comparison to control organoids (adjusted p-value: 0.0007, n=5) **(Fig. 3e)**. In contrast, the ADC of MMP-treated *LIS1^+/−^* organoids was not significantly different from controls. To support our findings, we performed three additional experiments that showed similar trends **(Supplementary Fig. S6a-d)**. These results demonstrate that *LIS1^+/−^* mutations contribute to increased organoid stiffness through altered ECM regulation, as evidenced by mass-spec data and the significant impact of MMP9 proteolysis on *LIS1^+/−^* CorticOs. DW-MRI further validates these findings by highlighting structural changes in ECM organization and stiffness in mutated and treated organoids.

### MMP9 treatment reversed numerous abnormal gene expression in *LIS1^+/−^* organoids

Finally, we enquired whether some of the DE genes in *LIS1^+/−^* will reverse in response to the treatment and its impact on the levels of stiffness and diffusion in mutated organoids shortly after the treatment. To emphasize potential rescue mechanisms, our attention was directed toward genes that exhibited differential expression between *LIS1^+/−^* and control groups prior to treatment but not afterward **(Fig. 4a, Supplementary Table S5)**. A heatmap of a subset of these genes is shown in **Fig. 4a**. These results indicate that at least part of the response is mediated through immediate (∼10 min) changes in gene expression. One possible mechanism for this quick recovery is a stiffness-sensitive expression of miRNAs.

**Fig. 4:**
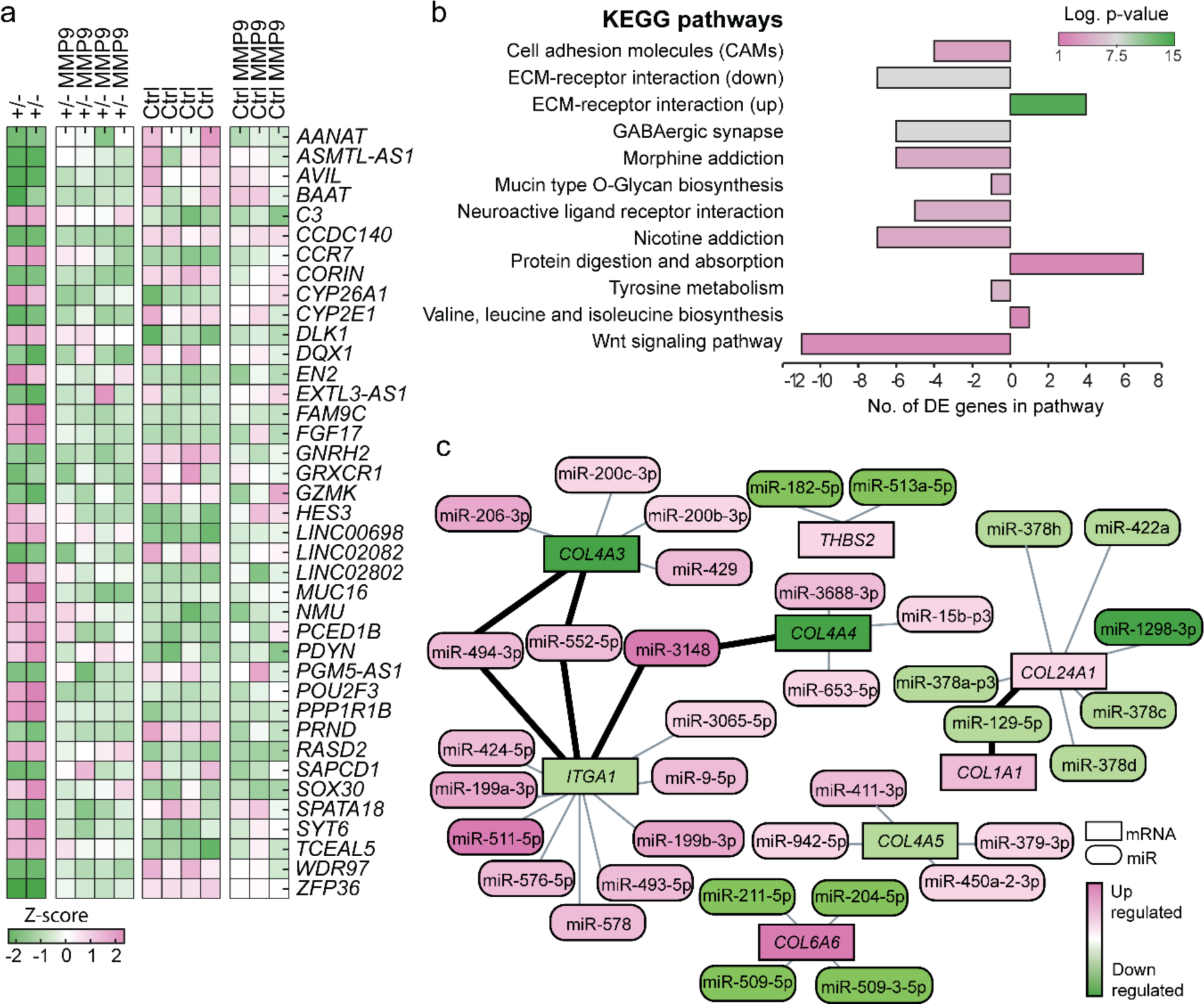
Rescued genes and inverse interaction between mRNA and miRNA in *LIS1^+/−^* organoids. **a**, A heatmap showing the top rescued genes and their expression before and after the MMP9 treatment. **b,** top KEGG pathways identified to be most influenced by the inverse relation between miRNA and mRNA expression. Negative numbers in the bar plot indicate that mRNA levels were reduced and that their targeting miRNAs increased. In contrast, positive values on the x-axis indicate an increase in mRNAs in the *LIS1*+/− samples while their targeting miRNAs were reduced compared to the control. **c,** Small RNA-seq integrated with target matrisome-related genes showing the opposite expression pattern of miRs and their predicted targeted mRNAs.

Previous papers reported that changes in the stiffness of a substrate used to grow cells affected the expression of a large group of micro-RNAs (miRs)XX. In addition, our group recently reported that *LIS1* is involved in regulating gene expression at several levels, including gene transcription, RNA splicing, and regulation of miRs. These changes were either dependent or independent of the Argonaute complex ^41^. Thus, we also conducted a small RNA-sequencing analysis which revealed 274 DE miRs **(Supplementary Fig. S7, Supplementary Table S6)**. These results suggest that at least some of the ECM expression abnormalities are linked to altered regulation of these genes’ expression by miRNA.

Using our existing database of the miRNA and mRNA expression of corticOs on day 105, we found that, indeed, the most significant interaction between DE mRNAs and miRNA in the *LIS1^+/−^* mutation was related to abnormal expression of genes associated with the ECM and the miRNA that are predicted to target and regulate their expression. Interestingly, when DE miRNA identified in small RNA seq were paired with DE mRNAs to identify regulatory mechanisms supported by both expression profiles, KEGG pathways enrichment analysis (http://microrna.gr/miRPathv3)^38, 42, 43^ pointed to the “ECM-receptor signaling” pathway (p < 0.0001) (**Fig. 4b**, **Supplementary Table S7**). Targeted genes highlighted by this miR-mRNA comparison included *COL4A3/4/5, COL5A3, ITGA1, COL1A1, THB2, COL24A1*, and *COL6A6* **(Fig. 4c)**. These results suggest that at least some of the ECM expression abnormalities are linked to altered regulation of these genes’ expression by miRNA.

### Collagen organization and a microstructure mechanical model

To further characterize the ECM abnormality, control, and mutated organoids were immunostained for several of the abnormally expressed structural ECM components in mutated and control organoids. While these immunostainings did not record the collagen enrichment, it did reveal substantial disorganization of the type 4 collagen and type 3 collagen fiber in 9 and 18 days old *LIS1*^+/−^ organoids. Whereas an apparent ring-like structure was observed in the control samples, the signal in the *LIS1^+/−^*samples was diffuse and punctate **(Fig. 5a-b)**.

**Fig. 5:**
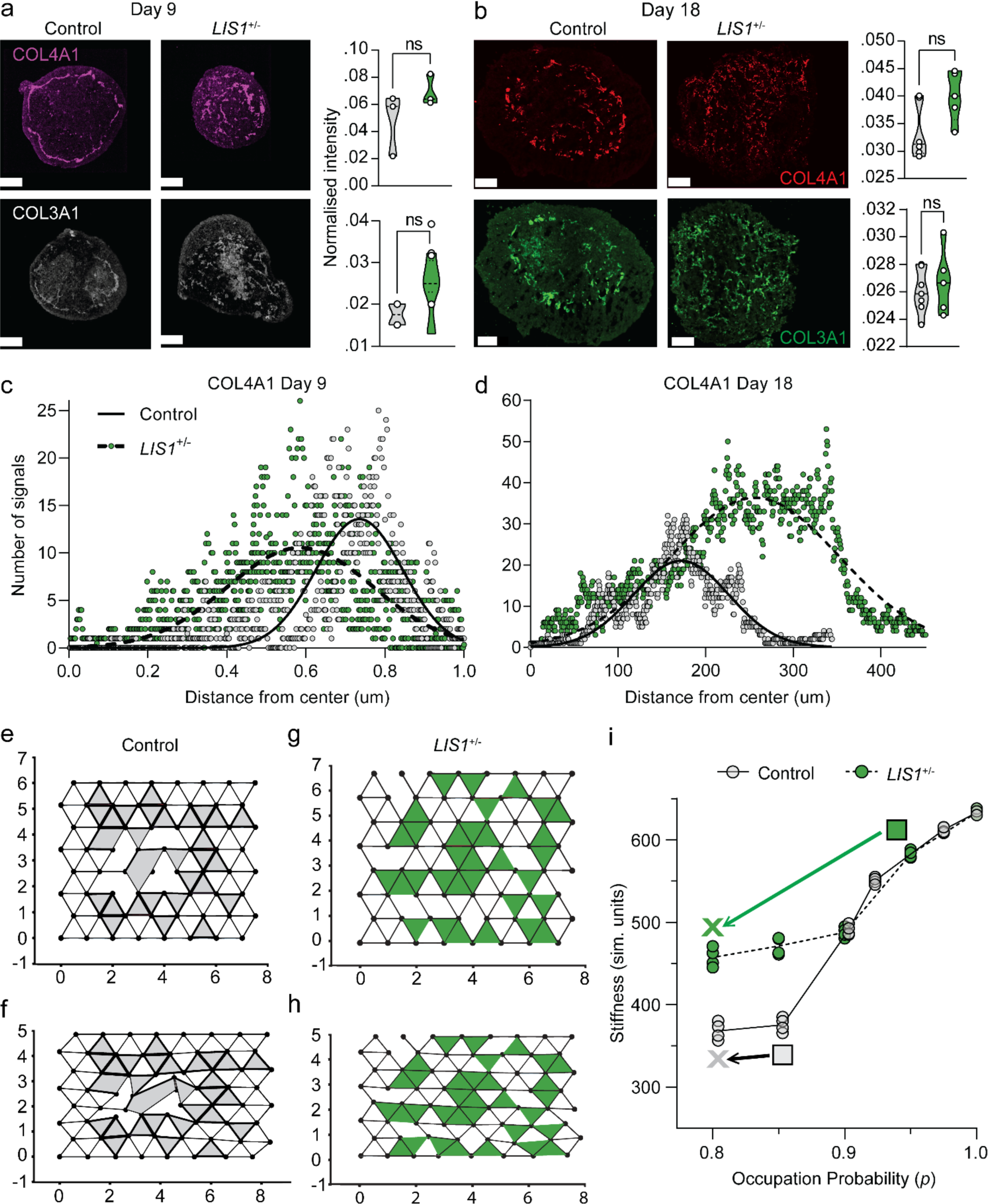
Collagen organization and a microstructure mechanical model. A-D. Immunostainings of COL4A1 and COL3A1 on **(a)** day 9 and **(b)** day 18, in control and *LIS1^+/−^* organoids with respective normalized intensity quantification of staining showing no difference between control and mutant organoids. Sholl analysis of COL4A1 signal in **(c)** day 9 and **(d)** day 18 control and *LIS1^+/−^* organoids represented as distribution from the center of the organoids highlighting the circular arrangement of collagen deposition in the control compared to an abnormal collagen distribution in the *LIS1*^+/−^ organoids. **e-f,** Simulation snapshots for 1% and 20% uniaxial compression strain in the control case. The control case is described by ECM removed within a localized circular region in the center of the system, as motivated by the ring of ECM. **g-h,** Simulation snapshots for 1% and 20% uniaxial compression strain in *LIS1*^+/−^ mutant case. The mutant case is described by randomly diluted ECM. **i,** Organoid stiffness (in simulations units) as a function of occupation probability *p*, which indicates the amount of ECM. Here, *N=64, N_c_=32*, and *R=10*.

Changes in the expression patterns were quantified using Sholl analysis, which demonstrated the signal clustered at a defined distance from the center in controls **(Fig. 5c-d)**. These findings suggest that the LIS1 protein is essential for the proper organization and localization of the ECM. We compared the *LIS1* mutant organoids with the control ones using additional immunostainings, and they had a greater number of SOX2^+^ and PAX6^+^ progenitors **(Supplementary Fig. S8a-b)**. However, they did not differ in the population of pVim^+^ radial glia or HOPX^+^ basal radial glia **(Supplementary. Fig. S8c-d)**. We did observe an increase in the number of pHH3^+^ mitotically active cells and the number of cells exiting the cell cycle (KI67^−^/EdU^+^), which may be attributed to the roles of LIS1 in cell cycle progression; no changes were noted in the population of KI67^+^ proliferating cells and EdU^+^ cells entering the S-phase. Furthermore, there were no significant differences in the number of apoptotic cells (Supplementary Fig. S9).

Finally, given the changes in the amount and spatial organization of the ECM between the control case and the *LIS1^+/−^* case, as well as the differences in stiffness between the two cases, we devised a more detailed mechanical model beyond the SLS model to help provide a quantitative link between the amount and organization of the ECM to brain stiffness. More specifically, we considered a computational model that includes both ECM fibers and cells and computed the stiffness of such a model. For simplicity, we investigated a two-dimensional model, or a cross-section, of the brain organoid. We will address this simplification in terms of what results we anticipate will carry over to three dimensions below.

The ECM was modelled as a triangular network of fibers, each consisting of both stretching energy and bending energy, with the latter encoding the semiflexibility of the collagen fibers ^54, 55^. In contrast, the cells are modelled as individual triangles randomly inserted in the lattice with some area stiffness ^56^ **(Fig. 5e-h)**. While the shapes of the cells in the computational model do not reflect the actual cellular shape, they do encode a key role in the mechanics of the cells. A line of edges on the lattice in one of the three principal directions represents a collagen fiber. To illustrate the disordered structure of the ECM, each edge in the triangular lattice was occupied independently and at random with occupation probability *p,* which also is a measure of the fraction of edges in the network. In other words, the larger the *p*, the more ECM and vice versa, and changing this *p* parameter encodes the amount of ECM.

Moreover, given the observation of more patterned ECM in the control, we also explored an ECM of a similar amount of ECM (similar *p*). However, the ECM is removed only within a localized region **(Fig. 5e-f)**. We then compared patterned ECM with randomized ECM in terms of influencing brain organoid stiffness. We note that such a model has already qualitatively captured nontrivial mechanical features of a fiber network with embedded particles, such as the phenomenon of compression stiffening ^56, 57^ (See Methods for a detailed description of the model and accompanying numerical calculations).

In control samples **(Fig.5e-f)**, as the brain organoid tissue is compressed with an increasing amount of uniaxial strain, the cells and fibers begin to distort; therefore, the energy of the tissue increases. We observed that the stiffness of the tissue, measured in simulation units, decreased as more ECM was removed from the localized circular region (**Fig. 5i),** as one goes from the grey box to the grey X). Notably, the decrease in stiffness is nonlinear, and the ECM provides the tissue with structural support and enhanced stiffness.

However, in the mutant case (**Fig. 5g-h**), where the ECM does not appear as organized and so the ECM randomly occupies edges on the network, we found that the tissue stiffness decreased nonlinearly as more ECM is removed (See **Fig. 5i)** as one goes from the green box to the green X). Interestingly, when the patterned ECM stiffness is compared to the random ECM stiffness for the same amount of ECM, the patterned ECM stiffness is not as large as the random ECM stiffness, at least for smaller amounts of ECM, i.e., smaller *p*. We argue that to compare the experiments to our computational model qualitatively, one can go from a smaller amount of ECM (lower *p*) on the patterned curve to a higher amount of ECM (higher *p*) on the random curve as indicated on (**Fig. 5i)** by going from the grey box to the green box. For instance, if we begin at *p*=0.85 fraction of edges occupied on the patterned ECM curve to 0.95 fraction of edges on the random ECM curve, we find a relative increase in the average stiffness of approximately 54%.

Moreover, this microstructure mechanical model qualitatively supports the results from the MMP9 treatment in which there was a greater decrease in organoid stiffness for the mutant case compared to the control. As decreasing the occupation probability also reduces the length of ECM fibers, the same trend is observed in our microstructure mechanical model as indicated by going from the green box to the green X versus going from the grey box to the grey X. Note that this treatment observation is more specific to the parameters at hand. These results were further tested under two additional cases of the parameter space, with more cells and with more stretchable fibers (**Supplementary Fig. S10a-b, respectively)**.

Overall, the computational model supports our findings that an upregulation of ECM enhances stiffness nonlinearly, which is not unexpected. However, our finding that the patterning of the same amount of ECM---diluted from a localized region as compared to randomly---affects organoid tissue stiffness is rather nontrivial in living tissue. The patterning creates weak spots, which, in turn, creates a weaker living material. Clearly, the computational model provides quantitative understanding of the trends observed in the experiments, including the MMP9 treatment findings. Finally, the computational model can provide predictions for brain organoid stiffness for decellularized material, which will presumably lead to compression softening ^56, 58^. The model can also provide predictions for the trend in any stiffness change for other mutated brain organoid should the organization of the ECM be altered in a different way from the mutant studied here.

## Discussion

Here, we studied the effects of LIS1 haploinsufficiency on ECM composition and organization and how these changes translate into modified physical properties of human brain-like organoids at different developmental time points. Our findings highlight LIS1 as an important regulator of ECM dynamics during human brain development. In both hippocampal and cortical organoid models, *LIS1^+/−^* mutations have been shown to significantly influence ECM composition and organization, leading to observable changes in tissue stiffness. Mass spectrometry analyses confirmed the enrichment of ECM-related proteins, particularly collagens, in *LIS1*^+/−^ organoids, indicating a mutation-driven alteration in ECM secretion and remodelling. These proteomic changes were complemented by biomechanical assessments using micropipette aspiration (MPA) rheology, revealing increased stiffness in *LIS1*^+/−^ organoids. Intriguingly, the application of MMP9, a zinc-dependent endopeptidase, on cortical organoids demonstrated a notable reduction in stiffness, especially in *LIS1*^+/−^ organoids, suggesting the reversibility of these changes through ECM proteolysis. Furthermore, Diffusion-Weighted Magnetic Resonance Imaging (DW-MRI) provided non-invasive insights into the structural organization alterations in these organoids, reinforcing the link between *LIS1* mutations, ECM dysregulation, and brain tissue mechanics. These results collectively highlight the critical impact of LIS1 on ECM regulation and its subsequent influence on brain development and structural integrity.

The ECM is not merely a passive substance between cells but an active source of signaling and regulating the stem-cell niche ^59–61^. It consists of molecules secreted by cells, serving as a structural scaffold while also harboring water reserves, growth factors, morphogens, and various bioactive molecules that engage neighboring cells and influence signaling processes ^62, 63^. Past studies identified that the ECM plays a pivotal role in the developing nervous system. For example, the synchronized movement of the neural plate and the mesoderm relies on the interplay between two ECM components, laminin and fibronectin. ^64^. Furthermore, the ECM is highly abundant within neuronal progenitors, and evidence shows that ECM enrichment is most pronounced in progenitors, which is uniquely prevalent in the developing human brain compared to the mouse brain ^22, 23^. *Ex vivo* experimental changes in the ECM affected the structure of human embryonic brain sections ^20, 21^. Our analyses revealed an abundance of matrisome-related proteins in *LIS1^+/−^* hippocampal and cortical organoids, whereas the composition of these two types of organoids exhibited distinctions. Previous studies have indicated that brain regions have ECM content variability ^65^. The observed proteomic changes in the corticOs partially correlated with corresponding transcriptomics data (mRNA and small-RNA), underscoring the roles of LIS1 in post-transcriptional regulation ^16, 24, 41^. LIS1 is an RNA-binding protein involved in post-transcriptional regulation and governs the physical properties of embryonic stem cells ^41^.

The *LIS1* mutation affected the physical properties of the brain-like organoids at multiple developmental stages; the *LIS1* mutant organoids were stiffer. The nuclei embedded within this more rigid tissue exhibited modulated parameters. *LIS1^+/−^*brain organoids exhibited increased levels of the nuclear lamina proteins, Lamin A/C, which are known to scale with increased stiffness ^39, 66–69^. In a correlative manner, the levels of DNA damage, indicated by γH2AX, decreased. The elevated levels of the ECM and increased rigidity are echoed in the Aperient Diffusion Coefficient (ADC) values. ADC values reflect the free water content of the tissue, which, in the case of brain-like tissue, is correlated with ECM ^70^. ADC values decrease in pathologies that involve cell swelling (edema) and can increase in chronic phases of stroke or other diseases involving necrotic cell death ^70, 71^. A monkey model employed to simulate a developmental brain disorder known as maternal immune activation (MIA), considered a model for autism spectrum disorders, displayed a significant elevation in the presence of extracellular free water within the gray matter of the cingulate cortex ^52^.

The physical measurements encompassing rheology and MRI appear to possess greater sensitivity compared to the protein-based assays involving mass spectrometry and specific collagen immunostainings. This heightened sensitivity is evident in the fact that we failed to observe a substantial rise in matrisome proteins through proteomics analysis or, in particular, collagens, through immunostaining for the early time points. However, immunostaining unveiled an unanticipated spatial arrangement of COL4A1 and COL3A1. While in control samples, they formed a circular structure, in *LIS1* mutant samples, they appeared scattered.

To consolidate our findings, we employed a computational model that takes into account the mechanics of cells, the ECM, and ECM organization. The control and the *LIS1* mutant differed in the amount of ECM and ECM organization and, hence, altered brain organoid stiffness, with the control case being less stiff than the mutant case in both the experiments and the computational model. Moreover, both the experiments and the computational model showed that effectively chopping up ECM enzymatically via MMP9 treatment led to a decrease in stiffness, with a pronounced decrease in the mutant case. Indeed, the mechanics and structure of brain organoids are intertwined as one helps determine the other. Computational models such as the one presented here, as well as other mechanical models ^72–76^ are, therefore, key to understanding the structure of brain organoids in both healthy and diseased states.

Moreover, multiscale, computational modeling tying the chromatin scale to the brain organoid, or tissue, scale to unravel the multiscale mechanics-structure relationship are on the horizon ^77^. Indeed, examining brain organoids from a materials point-of-view, as demonstrated here, provides novel perspectives about their structure and will help unravel the intricate mechanics-structure-function relationship in the developing brain and morphogenesis more generally. In summary, our study highlights the critical role of ECM composition and organization in human brain development and implicates the role of LIS1 in brain structure. Our findings suggest that targeting ECM components may be a potential therapeutic target in LIS1-related lissencephaly and possibly other brain malformations.

## Acknowledgments

We express our gratitude for the expert assistance provided by Drs. Alon Savidor, Yishai Levin, and Amir Prior from the de Botton Institute for Protein Profiling, in conducting proteomics experiments and analyzing the data. Orly Reiner is an incumbent of the Berstein-Mason professorial chair of Neurochemistry and the Head of the M. Judith Ruth Institute for Preclinical Brain Research. Our research has been supported by a research grant from Ethel Lena Levy, the Selsky Memory Research Project, the Gladys Monroy and Larry Marks Center for Brain Disorders, the Advantage Trust, the William and Joan Brodsky Foundation, and the Edward F. Anixter Family Foundation, the Nella and Leon Benoziyo Center for Neurological Diseases, the David and Fela Shapell Family Center for Genetic Disorders Research, the Abish-Frenkel RNA center, the Brenden-Mann Women’s Innovation Impact Fund, The Irving B. Harris Fund for New Directions in Brain Research, the Irving Bieber, M.D. and Toby Bieber, M.D. Memorial Research Fund, The Leff Family, Barbara & Roberto Kaminitz, Sergio & Sônia Lozinsky, Debbie Koren, Jack, and Lenore Lowenthal, and the Dears Foundation, a research grant from the Weizmann SABRA – Yeda-Sela – WRC Program, the Estate of Emile Mimran, and The Maurice and Vivienne Wohl Biology Endowment, the ISF grant (545/21), and the United States-Israel Binational Science Foundation (BSF; Grant No. 2017006).

The Alzheimer’s Association Grant AARG-NTF-21-849529 supported the development of the MRI procedure.

A.B. acknowledges the generous funding provided by BIRAX-Ageing that supported the design and operation of the micropipette aspiration apparatus.

JMS would like to acknowledge Mahesh Gandikota and Tao Zhang for the discussion and financial support from NSF-DMR-CMMT 2204312.

## Methods

### Cell lines

An NIH-approved human embryonic stem cell (ESC) line NIHhESC-10-0079, WIBR3 (W3), was used in this study. Isogenic mutant cell-line clones were previously generated by CRISPR-Cas9 mediated heterozygous deletion in the *LIS1* gene ^16^. Cell lines were checked for mycoplasma contamination regularly.

### Generation of hESC-derived corticOs and hippOs

hESCs were cultured in naïve media ^78^ until 70% confluency and then dissociated and aggregated in low adhesion wells (Day = 0). Aggregates were primed towards a neurogenic fate by application of the TGF-ß- and WNT-signaling inhibitors, SB-431542 and IWR-1, respectively (as specified in^45, 79^). These molecules facilitate neuroectoderm fate by inhibiting the mesodermal-promoting Nodal/Activin pathway ^80^; while preventing premature neuronal differentiation by inhibiting the WNT pathway via AXIN2 stabilization ^81, 82^ **(Fig.1A and 2A)**. On the 19^th^ day, aggregated neural precursors were introduced to conditions promoting either hippocampal or neocortical formation. In corticogenesis, the region of the dorsal pallium gives rise to the neocortex. In contrast, the medial pallium generates the hippocampus, positioned between the neocortex and the midline cortical hem. The cortical hem secretes dorsalising patterning morphogens such as WNTs and BMPs, which promote medial pallium formation. These also suppress the expression of FGF8 from the competent cortical primordium, which defines a more dorsal neocortical identity ^83^. Timely exposure (72h) to BMP4 and the WNT agonist CHIR-99021 was shown to be sufficient for inducing hippocampal fate by inhibiting GSK3ß and activating the WNT pathway^81, 82^, and was therefore used to generate hippocampal organoids (HippOs). For developing cortical organoids (CorticOs), aggregates were grown in a serum-free floating culture where they preferentially expressed FGF8. FGF8 then promotes self-organizing rostral-caudal polarity. After 35 days, CorticOs were treated with hLIF to induce bRG progenitor proliferation^36^.

### Immunohistochemistry

Ectoderm-like organoids were examined using immunohistochemistry 9 and 18 days after aggregation. Organoids were washed in PBS for 10 min at RT thrice, fixed for 30 min in 4% PFA, and washed again in PBS. Samples were dehydrated overnight at 4°C in 30% sucrose in PBS, embedded in OCT blocks and sliced into 16µm cryosections. Antigen retrieval was conducted in a water bath heated to 90°C in citrate buffer pH = 6 for 20 min and then chilled at RT. Tissues were permeabilized and blocked in a blocking solution (10% normal donkey serum [NDS] in PBST [0.1% Triton X-100]) for 3h and then incubated with primary antibodies for 48h at 4°C. After three washes, slides were presented with the secondary antibodies for 1:30h at RT at 1:200 concentration, after which they were incubated with 1:5000 4’,5-diamidino-2-phenylondole (DAPI) for 5 min. All the antibodies used in the study, their dilutions, and catalogue numbers are listed in method Error! Reference source not found..

### Image analysis and quantification

Immunostained slides were imaged with Oxford Dragonfly confocal microscope at 25X and stitched to get complete images of the whole organoid sections using the Fusion Shell software. The images were then analysed using the Spots Analysis of the Oxford Imaris software to determine the percentage of the total number of cells that expressed a particular cell-type specific marker and compare this parameter between control and mutant organoids. The total number of cells is considered the same as the total number of DAPI^+^ nuclei. Markers with nuclear expression like KI67, pHH3, EdU, SOX2, PAX6, HOPX and pVim were determined by setting up a colocalization filter under the Spots analysis.

For markers like COL3A1 and COL4A1, mean intensity of the signal in the total area of the organoid section was calculated in Fiji ImageJ software to compare their levels of expression. These values were then compared using statistical tests in GraphPad Prism to determine statistically significant differences, if any.

#### Sholl analysis quantification

The distribution of COL4A1 signal in day 9 and day 18 corticOs was determined using the Sholl analysis plugin (https://imagej.net/imagej-wiki-static/Sholl) in Fiji ImageJ. A straight line of uniform thickness was drawn diametrically through the organoid sections for the plugin to determine the centre of the section. Then the plugin calculates the number of signals encountered starting from the centre radially outwards in continuous concentric circles up to the edge of the sections. After multiple iterations, the software provides a distribution table, which was plotted in GraphPad Prism along with the nonlinear fit curve of distribution.

### Proteomics and Western Blot

Organoids were placed on ice and washed twice with 1X PBS for 5 min. In the third wash, organoids were placed in 100mM Tris-HCL, pH = 7.5. Proteins were extracted using lysis buffer (5% SDS 100mM Tris pH = 7.5). The lysates were transferred into soft tissue homogenizing CK14 tubes (Bertin Corp), placed in a homogenizer shaker at 400bps for 2 sec, and then put on ice for 20 min. Samples were centrifuged at 12,000g at 4⁰C for 20 min and then placed on ice. The uppermost, transparent liquid was transferred to a new Eppendorf tube and sent immediately to the De Botton Protein Profiling institute of the Nancy and Stephen Grand Israel National Center for Personalized Medicine, Weizmann Institute of Science for a ‘global quantification of protein’ mass spectrometry. This discovery experiment aimed to quantify as many proteins as possible using label-free-based methods. Samples included control and *LIS1^+/−^*mutated organoids hippOs and corticOs on days 70 and 105, respectively (N_hippOs_ = 3; n_hippOs_ = 5-8, N_coritcOs_ = 4; n_coritcOs_ = 6-8). Leftovers were aliquoted and used for western blot verifications.

### Micropipette aspiration

MPA was performed using a manometer setup as reported previously [CITE Wintner et al Av. Sci. 2021]. The organoids were placed immersed in media on a glass coverslip mounted on top of an inverted fluorescence microscope stage (Nikon Eclipse, Ti-U). The pipettes (Pipette brand and model) were washed in media with serum to lubricate the inner surface walls and decrease friction. Creep tests were performed over several seconds under 0.5-to-3 kPa suction-generated load consistent with the microenvironmental elasticity of healthy^47, 48^ and sclerotic brain tissues^45^.

For each organoid, the pipette inlet was brought into contact and creep test was performed by applying 1-to-3 kPa constant suction, which was measured relative to atmospheric pressure via a pressure transducer (Validyne, Northridge CA, USA). Timelapse imaging of the aspiration dynamics into the pipette was recorded over ∼ten sec and included ∼two sec before applying suction. We used pipettes with 0.15-to-0.5 mm inner radii to measure tissue level mechanics, thus probing integrated multicellular and ECM contributions. In specific, inner diameter 0.3 mm (B100-30-7.5HP, SUTTER INSTRUMENTS) was used on Day-9 and Day-18 cortocOs, 0.5 mm (B100-50-7.5HP, SUTTER INSTRUMENTS) for Day-35, and 1 mm (BF200-100-10, SUTTER INSTRUMENTS) for 70 days old coricOs.

This creep test response to applied load is characteristic of the minimal linear viscoelastic standard linear solid (SLS) model. In its Maxwell representation, it consists of an elastic element (spring *k_1_*) that is connected in parallel to a second elastic element (spring *k_2_*) positioned in series with a viscous element (dashpot *µ*) **(Fig. 1a’)**.

The effective time-dependent deformability of the organoids is given by the creep compliance function *J*(t), which we obtain over a range of small deformations using the half-space model relationship ^49^:

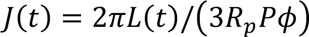

Here, the aspirated fraction *L*(t) length is scaled by the pipette inner radius *R_P_*. *P* is the applied pressure (relative to atmospheric pressure), and *Φ* = 2.1 is a geometrical factor that accounts for the pipette wall. To quantitatively characterize organoid mechanics, we fitted the SLS creep compliance function ^50^:

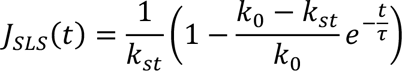

In this physically intuitive representation, the instantaneous stiffness *k_0_ = k_1_ + k_2_* measures the elastic resistance to abrupt impact. The steady-state stiffness that determines the long-term restoring forces is *k_st_ = k_1_*. The time scale for the transition from elastic stretching to steady-state deformation is estimated by *τ* = *μ* · (*k*_)_ + *k*_*_)⁄(*k*_)_*k*_*_). Satisfyingly, the R-square goodness of fit of all organoid measurements was high (typically > 0.98), confirming the choice of the minimal linear viscoelastic SLS model **(Supplementary Fig. S2b-f)**. Consistently, the goodness of fit of the mean creep compliance functions averaged across multiple organoids per condition were > 0.99 **(Fig. 1b-e)**.

### ELISA activity assay

The MMP9 catalytic domains were incubated in a 96-well plate with different concentrations of Mca-PLGL-Dpa-AR-NH2 Fluorogenic MMP Substrate (ES001, R&D SYSTEMS), a substrate that is cleaved by the MMP9. The peptide substrate contains a highly fluorescent 7-methoxy coumarin group that is quenched when cleaved. Active peptidases that can cleave an amide bond between the fluorescent group and the quencher group cause an increase in fluorescence, which can confirm their function. Each well was supplemented with 20nM of either MMP9, a TNC digestion buffer (50mM TRIS-HCl pH = 7.5, 150mM NaCl, 5mM CaCl2, and 0.05% Brij), and the substrate at a dilution series (1-100uM). Two control conditions were ‘TNC-only’ and a 10uM ‘TNC + substrate’ condition. While in 37⁰C, fluorescence was captured by a plate reader (Synergy H1) every 50 seconds, for an overall duration of 40 minutes **(Supplementary Fig. S2)**.

### MMP9 catalytic domain treatment

On day 18, control and *LIS1^+/−^* organoids were treated with 500 µM of MMP9 catalytic domain diluted in the organoids’ original media and incubated for 10 min in 37⁰C, 5% CO_2_. The treated media was then replaced with fresh 100 µl of ‘SA1 media’, and plates were held in the incubator until the organoids were tested.

### Diffusion MRI – data analysis

The Diffusion Weighted Imaging (DWI) MRI dataset was collected to assess the differences between brain-organoid groups. Based on the collected dataset, the Aperient Diffusion Coefficient (ADC) maps were calculated using a mono-exponential fit. The images with the highest b-value (the degree of diffusion weighting) were used to segment and identify the voxels of the brain organoid tissue. **Supplementary Fig. S5a** shows the ADC maps and the contour of the segmented voxels. A normalized distribution of the ADC values in the identified voxels for each type, consisting of 100 bins, was defined. A denoising of the distribution profile was then performed, removing high-frequency components using FT. The distribution of the diffusion in the brain organoids tissue results in an asymmetric profile; therefore, the maximal likelihood position was defined as a centre of points with 2/3I_max_. intensity (Imax was found based on the denoised distribution and the two points with 2/3I_max_ intensity from both sides of the distribution by interpolation). The same steps were repeated for each organoid type and scan. **Supplementary Fig. S5a-c** shows the ADC values distributions and the estimated maximal likelihood positions. Note that the ADC values increased as a function of the repeated scans. This can be due to the long scan duration through which the organoids were out of their regular environment. However, the deviation of LIS1^+/−^ compared to Control was preserved over the scan duration. Three additional experiments were performed with different subsets of the organoid types. **Supplementary Fig. S5d** shows a similar observed trend in the repeated experiments. The ages of the organoids in the different experiments were: experiment 1, 19 days; experiment 2 & 3, 18 days; experiment 4, 16 days. The lower ADC values in Exp.#4 can be due to different scan parameters (with a slice thickness of 100 μ compared to 200 μ; see the scan parameters summarized below.

MRI scanner details: Horizontal Bruker Biospec 15.2-T USR preclinical MRI scanner with an Avance IIIHD console.

Container used for the MRI scans: In experiments #1-3, the organoids were placed in a container with separated wells for each type, allowing for scanning all types in one scan. Exp. #4 was scanned with a single Eppendorf in which the organoids of one kind were placed and scanned, and then the scan was repeated with the second type. The medium used was PBS.

RF coil details: 1H MRI CryoProbe 2-element array kit for the mouse head.

The DWI scan parameters:

- Exp.#1: TR/TE 400/13.7 ms, FOV 12×12 mm^2^, in-plane resolution 100×100 μ^2^, slice thickness 200 μ, 20 slices, ten averages, scan duration 48 minutes. The scan was repeated five times. The scan included six b-values - 0, 200, 400, 600, 1000, 1200 s/mm^2^.
- Exp. #2: TR/TE 400/13.7 ms, FOV 18×12 mm^2^, in-plane resolution 100×100 μ^2^, slice thickness 200 μ, nine slices, ten averages, scan duration 48 minutes. The scan included six b-values - 0, 200, 400, 600, 1000, 1200 s/mm^2^.
- Exp.#3: TR/TE 400/13.7 ms, FOV 12×12 mm^2^, in-plane resolution 100×100 μ^2^, slice thickness 200 μ, 20 slices, 20 averages, scan duration 1h36m0s0ms. The scan included six b-values - 0, 200, 400, 600, 1000, 1200 s/mm^2^.
- Exp.#4: TR/TE 300/12.7 ms, FOV 12.6×2.4 mm^2^, in-plane resolution 100×100 μ^2^, slice thickness 100 μ, 60 slices (3D acquisition), one averages, scan duration 1h12m0s0ms. The scan included four b-values - 0, 200,600,1000 s/mm^2^.

#### RNA and miRNA sequencing

Total RNA from 105 days old CorticOs (N = 4; n = 6-8 per genotype) was extracted using the Direct-zol RNA Miniprep Plus kit (Zymo Research) following the manufacturer’s protocols. RNA concentration and integrity were measured using Nanodrop (Thermo Scientific). Samples were then sent to the DNA Link Sequencing Lab in S. Korea, where RNA-seq libraries were prepared using TrueSeq standard mRNA library kit or small RNA-seq libraries. Libraries were sequenced in NovaSeq6000. The raw data was processed using the Weizmann Institute bioinformatics pipeline. DE genes were analysed using Metascape ^84^ and gene analytics. The processed RNA-seq databases are available at the GEO accession: GSE228926 (Go to https://www.ncbi.nlm.nih.gov/geo/query/acc.cgi?acc=GSE228926 and enter token **avkdiyuotvavvoj** into the box).

### Computational Model

The energy functional for the mechanics of the microstructure of a brain organoid with N nodes and N_C_ number of cells:

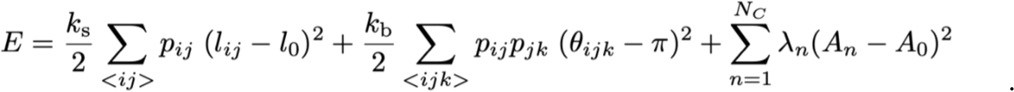

Here, *k_s_* denotes the individual ECM fiber stretching stiffness, while *k_b_* denotes the individual ECM fiber bending stiffness to give to a persistence length. Moreover, *p_ij_* represents the edge occupation probability, or the fraction of edges in the network that are occupied. We set *p_ij_* = *p* for convenience in the main body of manuscript. As *l_ij_* quantifies an edge length in the network, the first term in the equation above captures the stretching energy. As *8_ijk_* denotes the angle between two edges, the second sum in the equation above quantifies the semi-flexibility of the fiber network, while the third term encodes the bulk stiffness of *N_c_* cells each area *A_n_* and stiffness **λ*_n_=*λ*q_n_*, with *q_n_* either zero or one as the number of cells is set and then the triangles in the lattice are randomly occupied with cells. Therefore, *λ* quantifies the individual bulk cell stiffness, which is, for simplicity, the same for all cells. Note that *k*_s_/(*k*_b_*l*_0_^2^) = *R* and is the dimensionless ratio comparing the individual ECM fiber stretching stiffness to bending stiffness.

To measure the stiffness *K* of this composite system, once the model parameters are chosen, we impose periodic boundary in the horizontal direction and cut, delete the periodic edges in the vertical direction, and then compress the system in the vertical direction by applying strain *γ.* We then compute the energy *E* of the system, the first derivative of the energy to obtain the strain and, finally, the second derivative of the energy of the system to compute the material stiffness *K,* which includes a *1/A_0_* prefactor. We make such measurements for different amounts of randomly positioned ECM (mutant) as well as for patterned ECM (control) via the removal of ECM from localized circular region near the center.

The code related to this manuscript is deposited in: https://github.com/jenschwarz/CellsFibers2DMechanicalModel (at the present time private and public upon acceptance).

### Table of antibodies

| Application | Target | Species | Dilution | Company | Catalog number |
| --- | --- | --- | --- | --- | --- |
| IHC | Collagen type 3 α-chain 1 | Rabbit | 1:200 | Abcam | ab7778 |
| IHC | Collagen type 4 α-chain 1 | Rabbit | 1:500 | Abcam | ab236640 |
| IHC | FAM107A | Rabbit | 1:250 | Sigma | HPA055888 |
| IHC | GFAP | Rabbit | 1:2000 | DAKO | Z033401-2 |
| IHC | HOPX | Rabbit | 1:250 | Sigma | HPA030180 |
| IHC | HOPX | Mouse | 1:50 | Santa Cruz | 398703 |
| IHC | LEF1 | Rabbit | 1:200 | Cell signalling | 2230 |
| IHC | NeuN | Mouse | 1:200 | Millipore | MAB377 |
| IHC | PAX6 | Chicken | 1:250 | DSHB | AB528427 |
| IHC | PAX6 | Rabbit | 1:100 | BioLegend | 901301 |
| IHC | pVIM | Mouse | 1:100 | MBL | D076-3 |
| IHC | SOX2 | Mouse | 1:300 | Santa Cruz | sc-365823 |
| IHC | TUJ1 | Mouse | 1:200 | Convance | Ab18207 |
| IHC | ZBTB20 | Rabbit | 1:200 | Sigma | HPA016815 |
| IHC | MAP2 | Mouse | 1:200 | Sigma | M9940 |
| IHC | KI67 | Mouse | 1:400 | BD Pharmingen | 550609 |
| IHC | pHH3 | Rabbit | 1:200 | MilliporeSigma | 06-570 |
| IHC | cleaved Caspase3 | Rabbit | 1:200 | Cell signaling | 9661 |
| WB | LMNB1 | Rabbit | 1:10,000 | Abcam | ab16048 |
| WB | LMNA/C | Mouse | 1:5,000 | Cell signalling | 4777 |
| WB | γH2AX | Rabbit | 1:50 | R&D systems | AF2288 |
| WB | GAPDH | Rabbit | 1:10,000 | Cell Signalling | 2118 |
| WB | ACTIN | Mouse | 1:10,000 | Sigma | a5441 |

## Notes

### Competing Interest Statement

The authors have declared no competing interest.

