## Supplementary Figures for "Altered Extracellular Matrix Structure and Elevated Stiffness in a Brain Organoid Model for Disease"

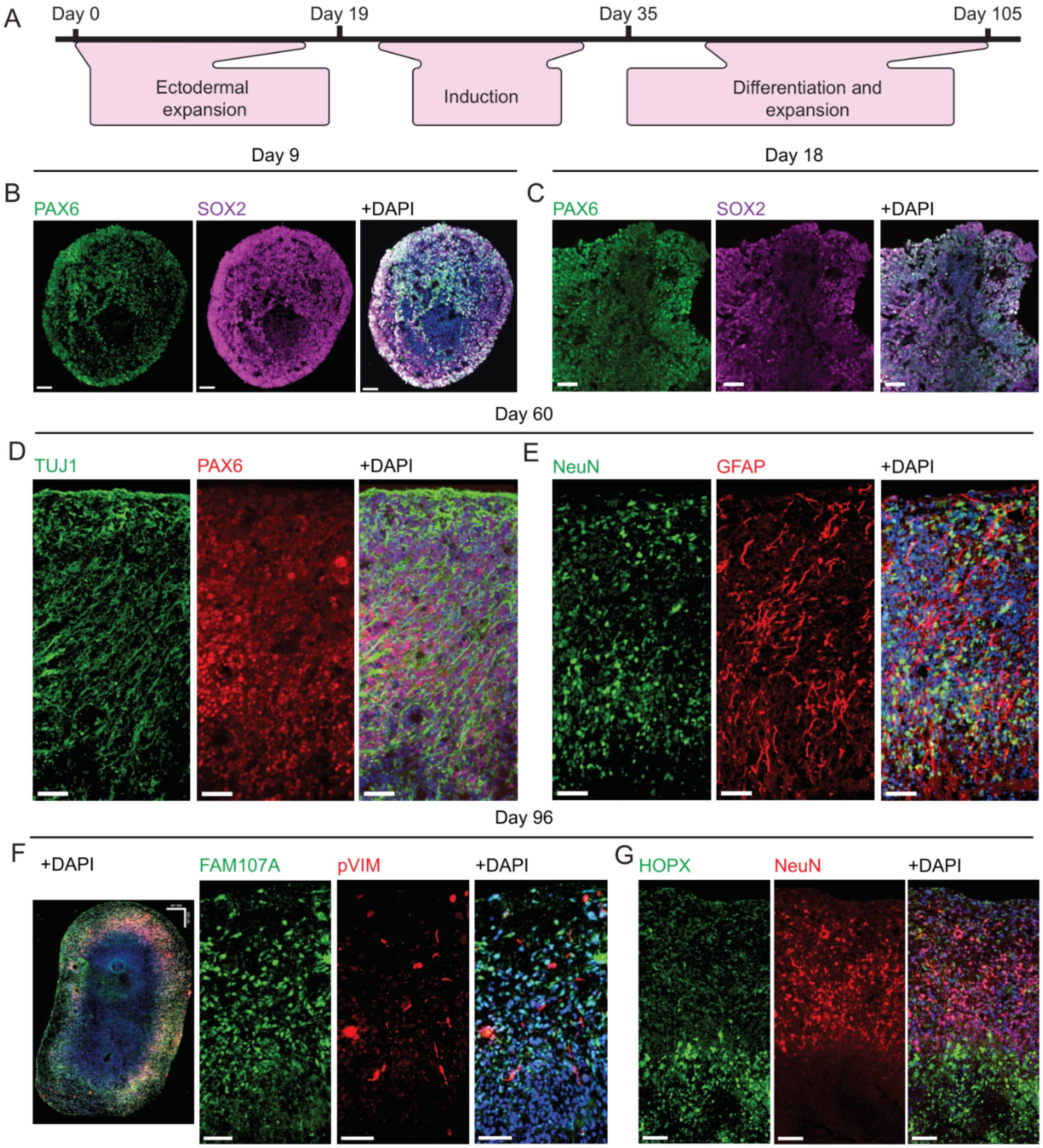


**Supplementary Fig. S1: Cortical Organoids (CorticOs). a,** Timeline of corticOs. **b-d,** Proliferative markers SOX2 and PAX6 in ectodermal-like organoids after 9 days (**b**) as well as on day 18 (**c**). **d-e,** PAX6$^+$ cells also appeared on day 60, along with neuronal processes marker TUJ1 (**d**), NeuN$^+$ neurons, and GFAP$^+$ astrocytes (**e**). **f-g,** On day 96, FAM107$^+$ and pVIM$^+$ (**f**), HOPX$^+$ basal radial glia cells were expressed, and NeuN$^+$ neurons (**g**). All figures have a scale bar of 50 µm.

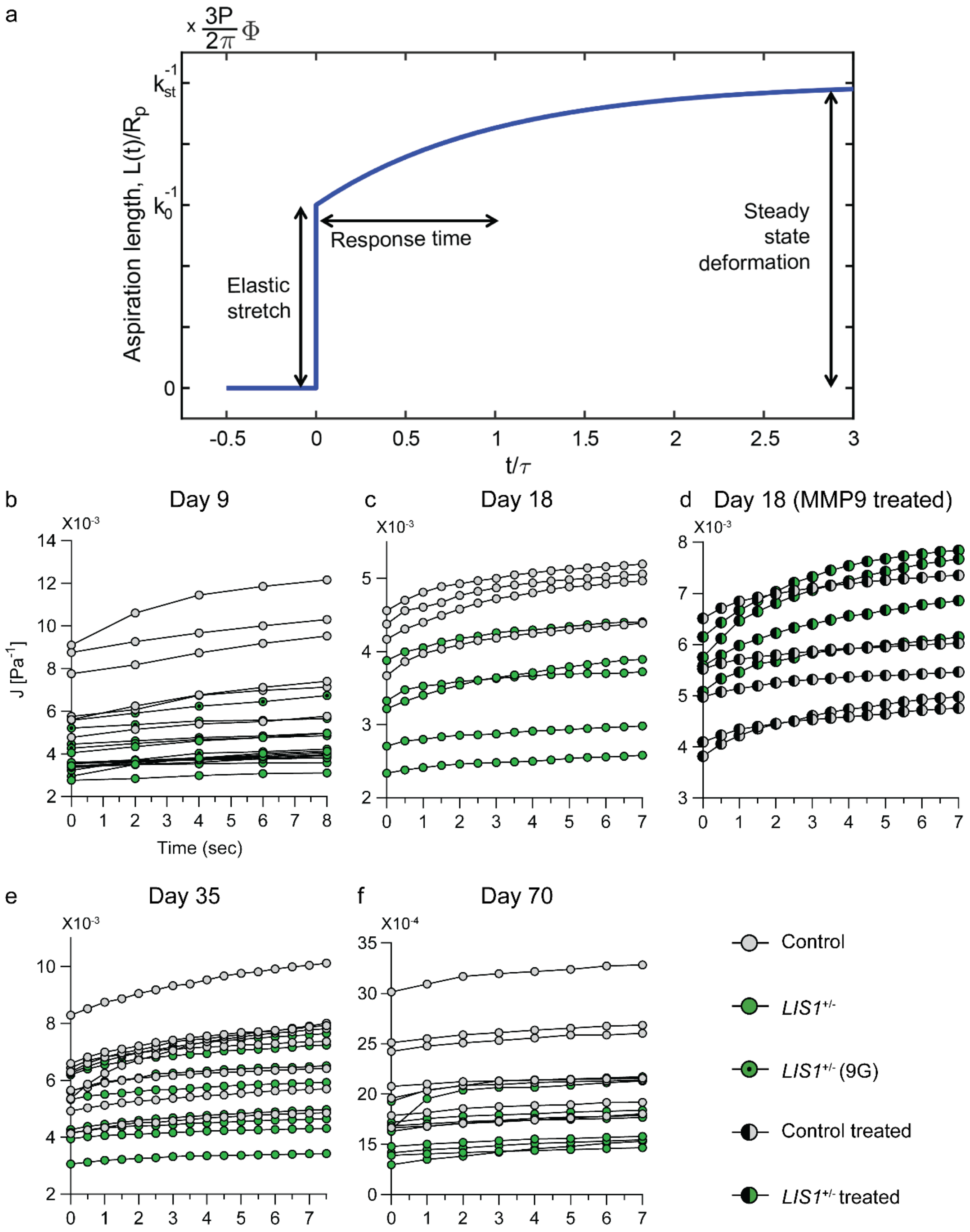


**Supplementary Fig. S2: Creep test MPA rheology. a,** The aspiration length $L(t)$ of an SLS material in a creep test is presented. The material stretches elastically to an aspirated length that is proportional to ${k_0}^{-1}$, then aspiration continuous over a length scale $\tau$ that marks viscoelastic deformation before approaching a steady-state deformation with aspiration length that is proportional to ${k_{st}}^{-1}$. $k_0 = k_1 + k_2$, $k_{st} = k_1$ and $\tau = \mu \cdot (k_1 + k_2)/(k_1 k_2)$ are the viscoelastic elements of the SLS material (Fig. 4A). **b-f,** Creep test measurements of Day-9, Day-18, Day-35 and Day-70 organoids per the specified treatments. One measurement was performed for each organoid. Unless specified otherwise, "treated" refers to X.

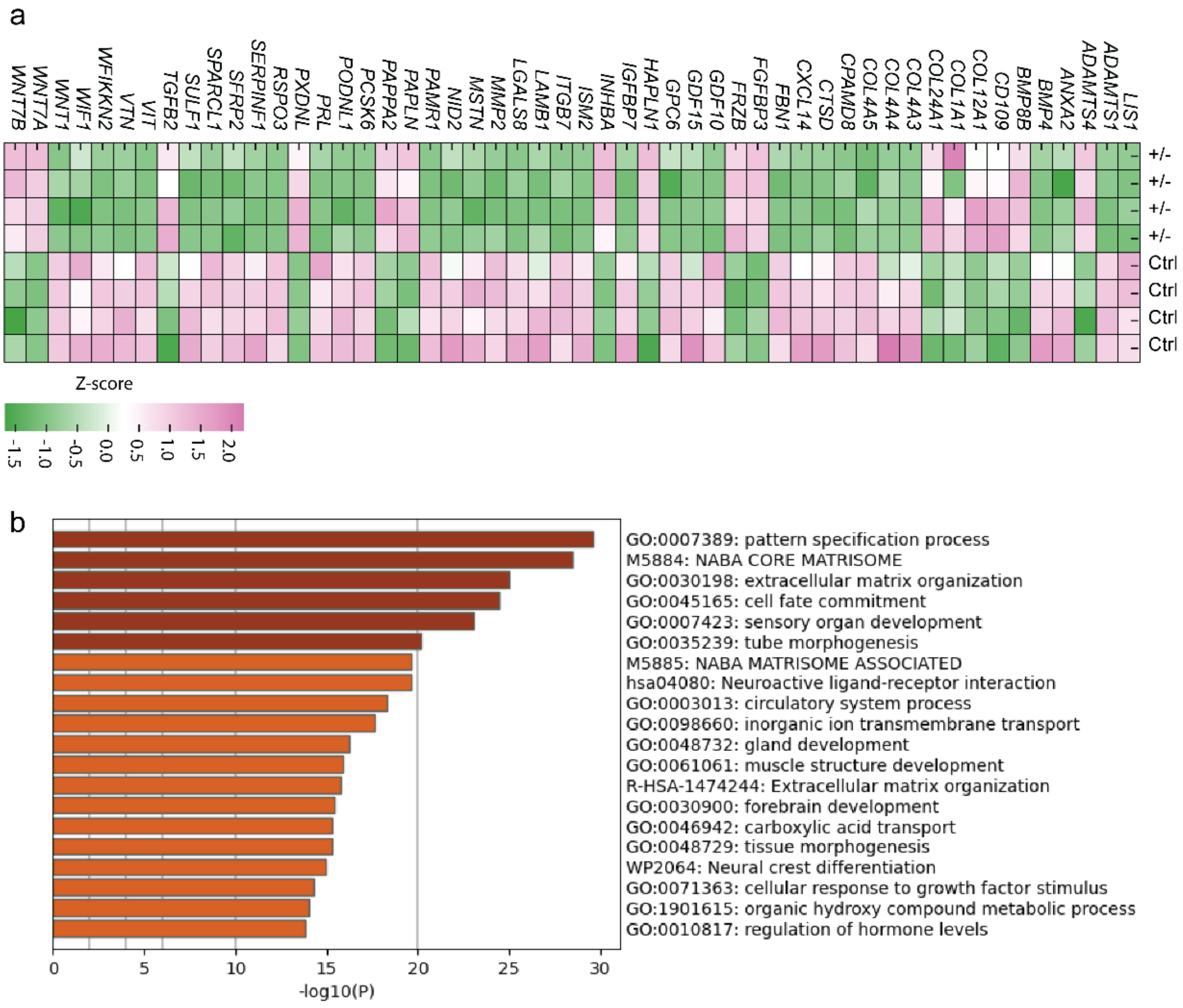


**Supplementary Fig. S3: RNA-seq of 105 days old corticOs reveal altered ECM expression. a,** Heatmap of top DE mRNAs between control and *LIS1*$^{+/-}$ mutant organoids, after logarithmic transformation of counts and z-score normalisation. **b,** Metascape analysis of 2064 genes that are DE between control and *LIS1*$^{+/-}$mutant organoids

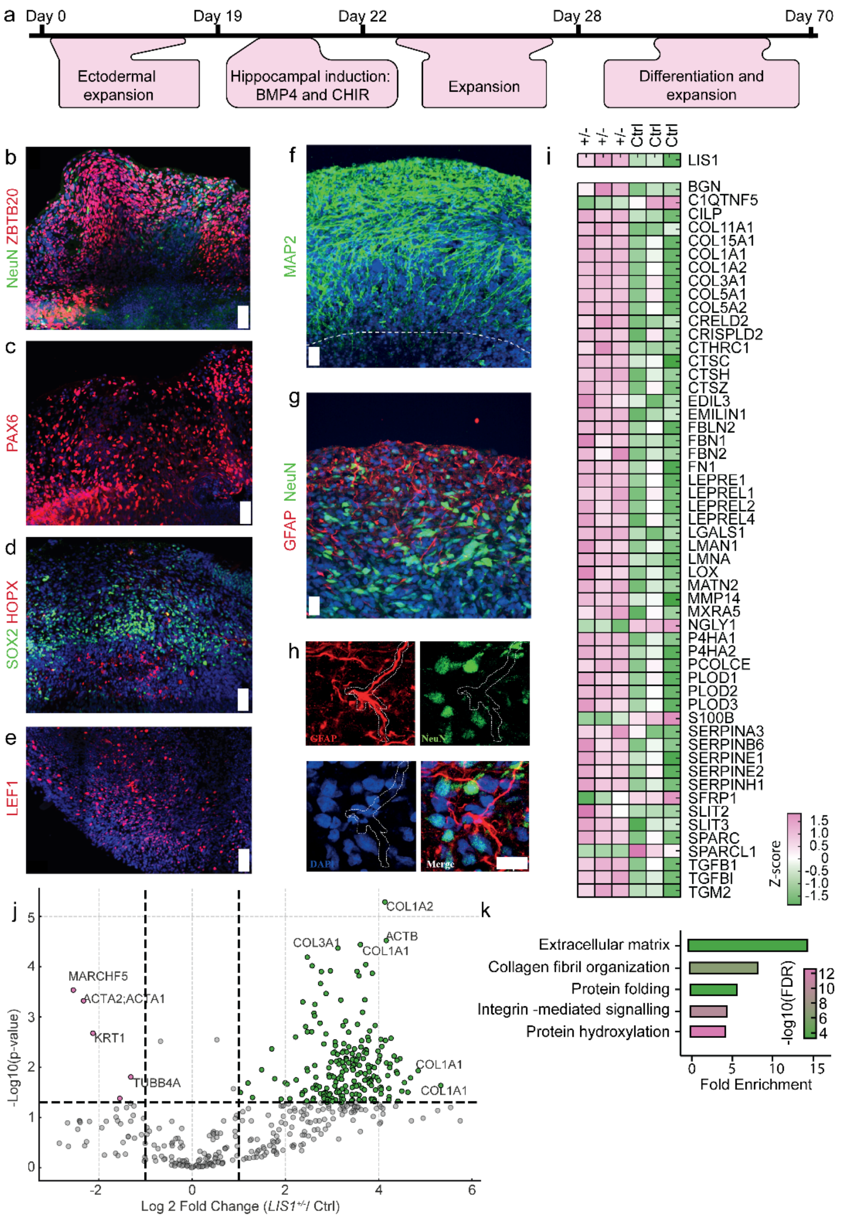


**Supplementary Fig. 4: Hippocampal organoids characterization and comparative proteomics of *LIS1*$^{+/-}$ and control samples. a,** Timeline of hippocampal organoid cultures. **b-e,** After seventy days, hippOs expressed the post-mitotic neurons marker NeuN and the dentate gyrus marker ZBTB20 (**b**), the progenitor markers PAX6 (**c**) HOPX, and SOX2 (**d**), and the transcription factor LEF1. **f-h**, MAP2$^{+}$ neuronal processes markers, (**f**) GFAP$^{+}$ astrocytes, and NeuN$^{+}$ neuronal cells (**g**), as well as a close-up visualization of these markers. **i**, Heatmap of DE matrisomal proteins in *LIS1*$^{+/-}$ and control hippocampal organoids. Cell color expresses normalized reads following logarithmic transformation and Z-score normalization. **j,** Volcano plot of hydroxylation in the MS peptide reads. Points marked in green and pink show the top enriched PTM targets in the *LIS1*$^{+/-}$ and control organoids, respectively. **k,** False discovery rate of DE proteins following a WebGestalt Gene Set Enrichment Analysis (GSEA) analysis [78]. Scale bars in **b-e** demonstrate 50 μm and 10 μm in **f-h**.

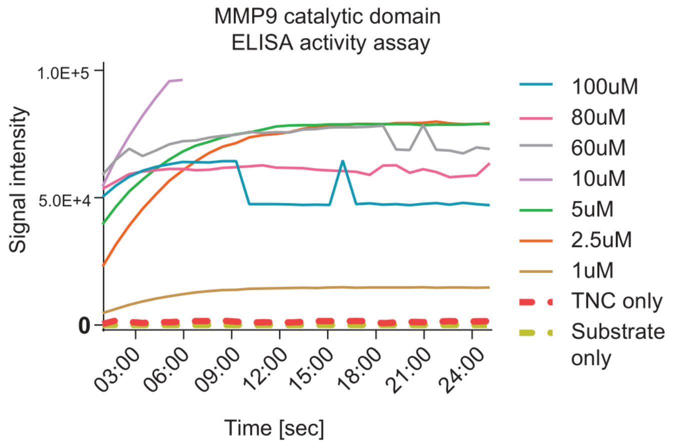


**Supplementary Fig. S5: MMP9 catalytic domain ELISA activity assay.** Fluorescence signal intensity at uniform time intervals for two control conditions - 'TNC-only' and 'TNC + substrate' and experimental conditions with varying concentrations of MMP9 catalytic domain (from 1 μM to 100 μM).

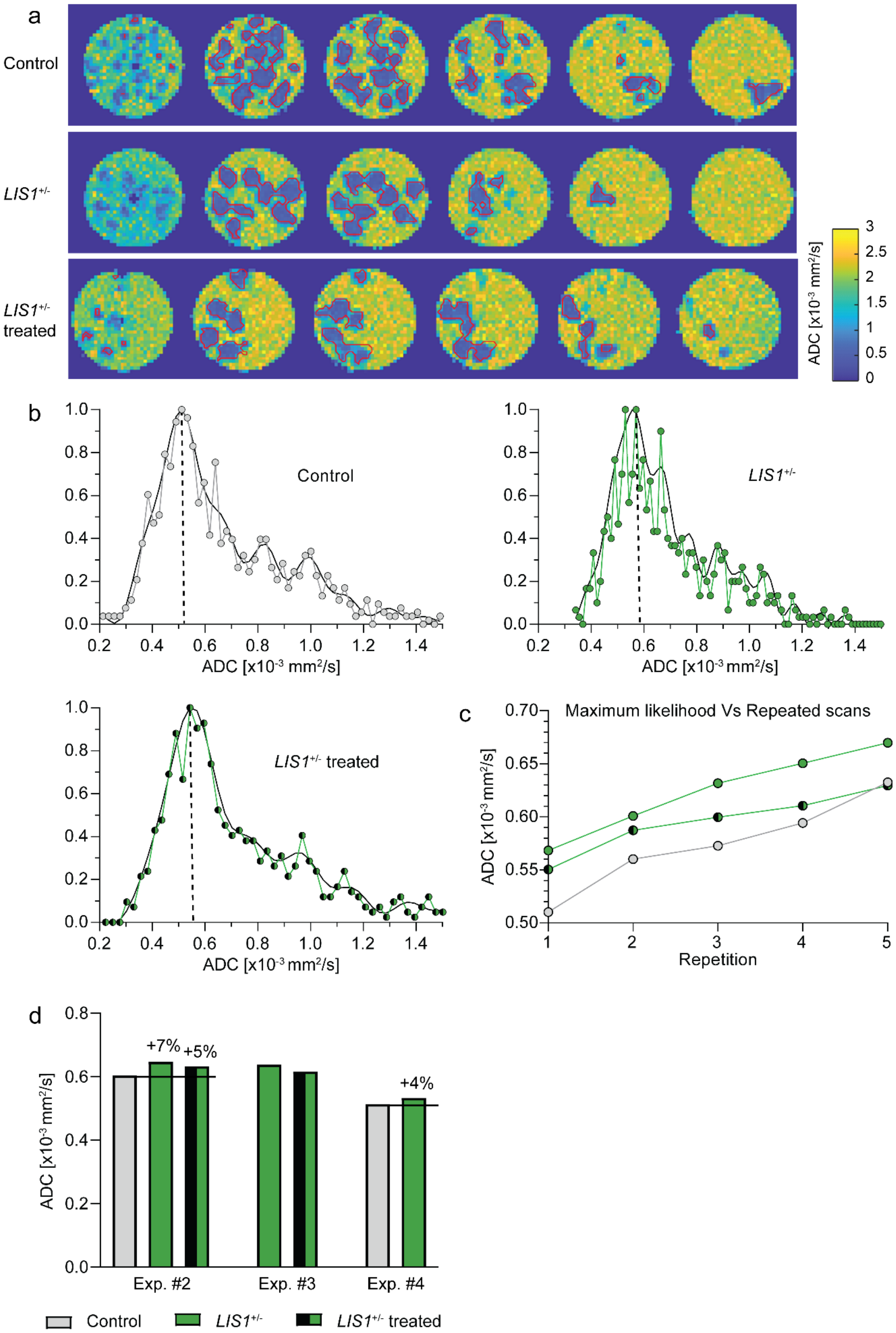


**Supplementary Fig. S6: MRI additional information. a,** ADC maps including an overlay of identifying contours of control, *LIS1*$^{+/-}$ and MMP9-treated *LIS1*$^{+/-}$ CorticOs in Exp.#1 (from top to bottom). Number of voxels for each group was 942, 618,627, respectively. **b,** ADC distribution and the estimated maximum likelihood position from a single scan in Exp.#1 (shown for all three groups). **c,** Estimated maximal likelihood position of the ADC as function of the repeated scans in Exp.#1. **d,** Estimated ADC maximum likelihood positions in three additional experiments (Exp.#2-4). An average number of voxels for each experiment was: 966, 3925, 3176.

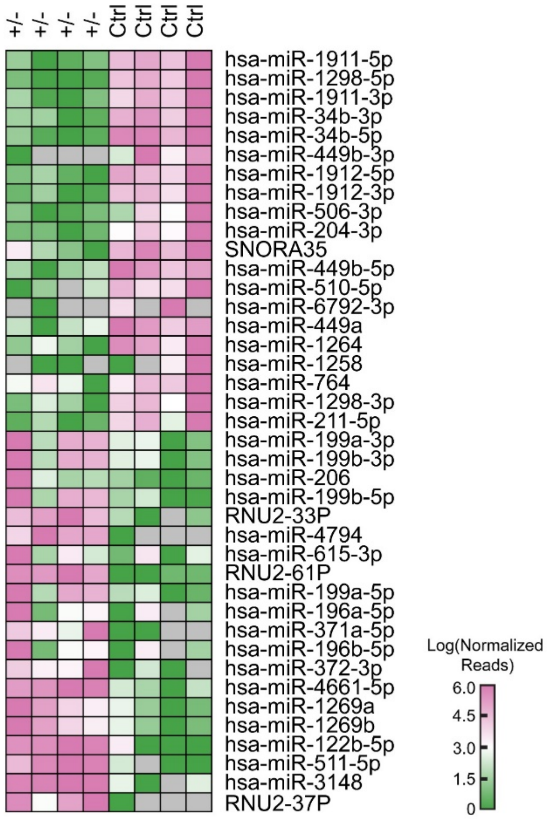


**Supplementary Fig. S7: miRNA-seq of 105 days old corticOs.** Heatmap of top miRNAs DE between control and *LIS1*$^{+/-}$ organoids. The scale for the heatmap represents normalised reads on logarithmic scale.

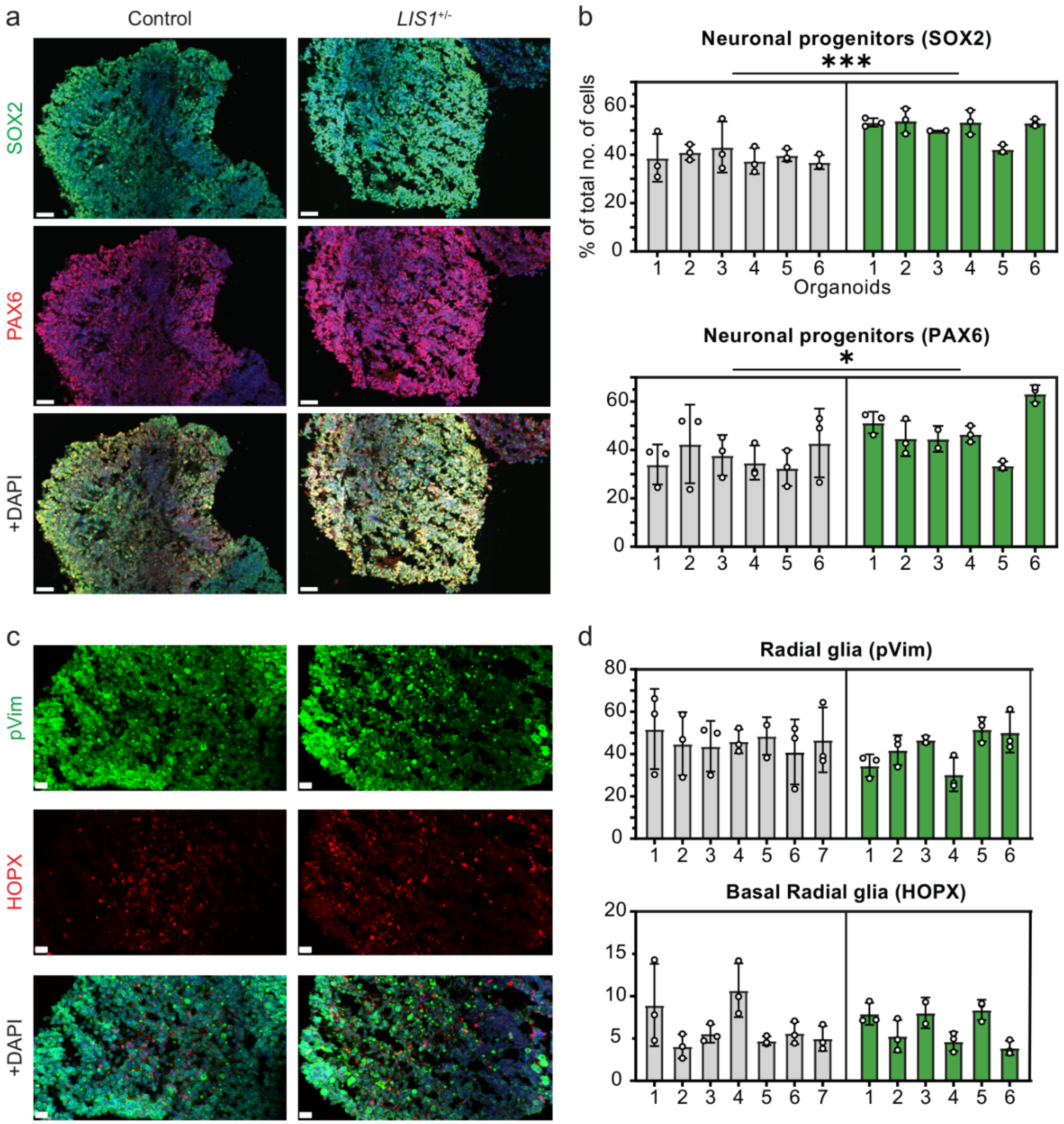


**Supplementary Fig. S8: Progenitors and neurons in CorticOs.** Immunohistochemistry of 18 days old control and *LIS1*$^{+/-}$ organoids showing **(a)** neural progenitor markers SOX2 and PAX6, and **(c)** basal radial glia marker pVim and HOPX, **b,** Percentage of the total number of cells that are progenitors (SOX2$^{+}$, PAX6$^{+}$), and **(d)** pVim$^{+}$ and HOPX$^{+}$ cells on day 18 (Nested One-Way ANOVA, n = 7, α = 0.05). Images scale bar represents 50 µm.

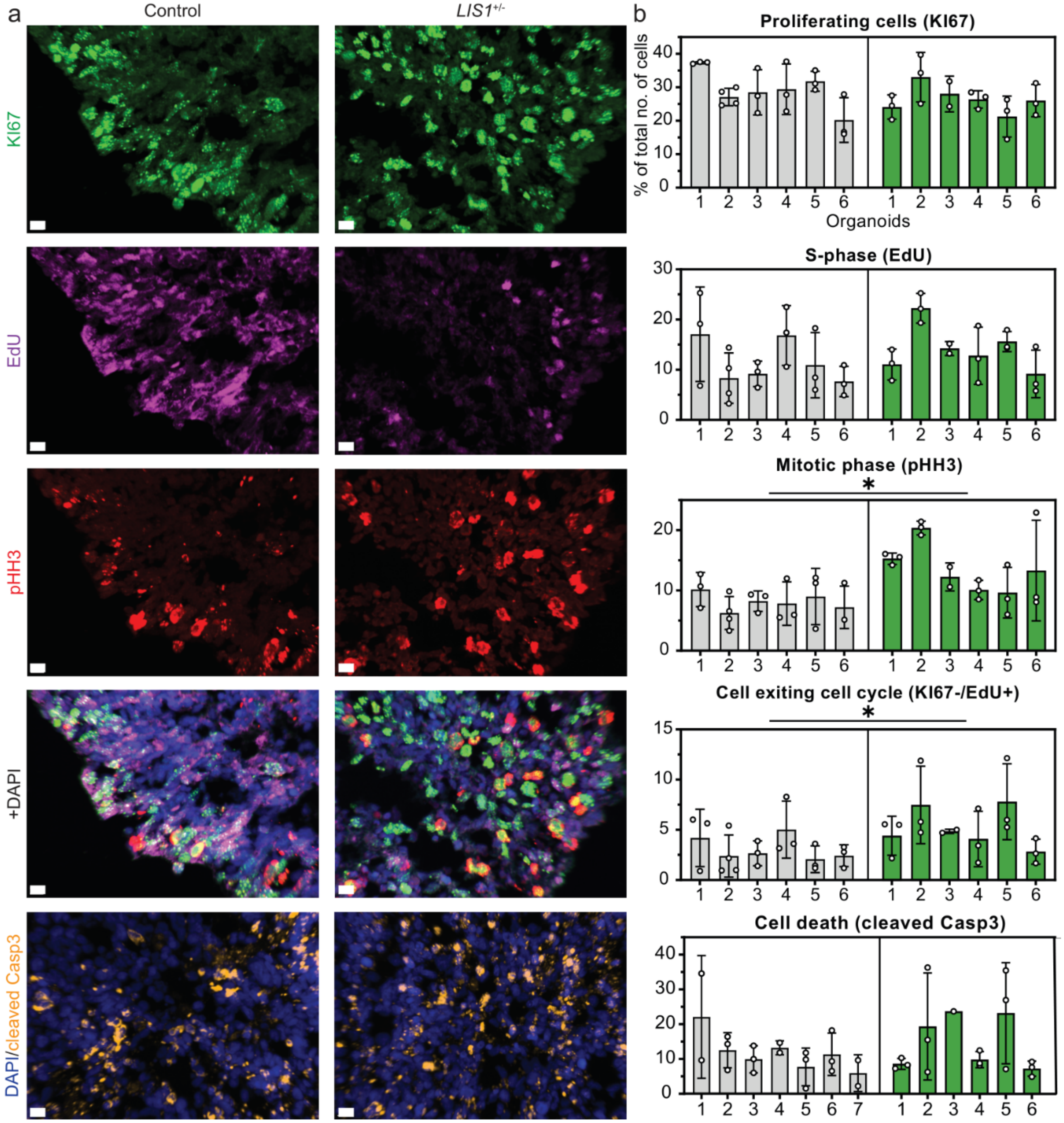


**Supplementary Fig. S9**: **Proliferation and cell death in CorticOs.** a, Immunohistochemistry of 18 days old control and LIS1$^{+/-}$ organoids showing the proliferation markers – KI67, EdU, pHH3, and cell death marker cleaved Casp3). b, Percentage of the total number of cells that are proliferative (KI67$^{+}$), in initial S-phase (EdU$^{+}$), mitotically active (pHH3$^{+}$) and undergoing cell death (cleaved Casp3$^{+}$) respectively, in day 18 control and LIS1$^{+/-}$ organoids (Nested One-Way ANOVA, n = 7, α = 0.05. Scale bar of images represents 10 μm.

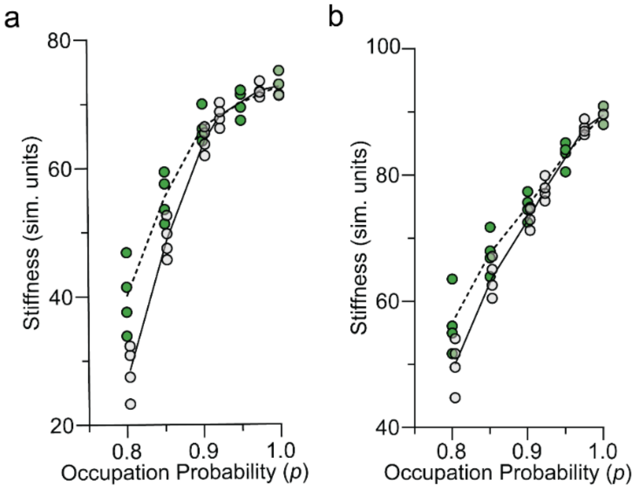


**Supplementary Fig. S10: Computational model scenarios of stiffness prediction. a,** Same plot as in **Fig. 7i** but with $R=1$by decreasing $k_s$ by a factor of 10, resulting in a smaller stiffness. **b,** Same plot as in the previous supplement figure with an increased number of cells ($N_C=42$). With the decreased $R$, we did not find as significant a difference in stiffness between the control case and the mutant case as in (**Fig. 7i)**. However, the trend of a larger $R$ (**Fig. 7i**) is more in line with stiffer collagen fibers. Moreover, by increasing the number of cells the mechanics is becoming less dominated by the fibers and more by the cells, the former of which is organized similarly between the two cases, hence, more similar stiffnesses even for smaller occupation probabilities.
